# Conformation-gated binding drives negative cooperativity in ATP:cob(I)alamin Adenosyltransferase for optimized cobalamin handling

**DOI:** 10.1101/2025.01.07.631765

**Authors:** Guangjie Yan, Manhua Pan, Aaron M. Keller, Ace George Santiago, Michael Lofgren, Ruma Banerjee, Peng Chen, Tai-Yen Chen

**Author notes:** Tai-Yen Chen. **Email:**. Peng Chen. **Email:**. **Author Contributions:** P.C. and T.-Y.C. designed research and directed the earlier (P.C.) and later (T.-Y.C.) phase of research; M.L. and R.B. designed plasmids for protein expression, and provided some of the purified recombinant protein used in this research; G.Y., M.P., A.M.K., A.G.S. and M.L. performed experiments; G.Y. and M.P. contributed new analytical tools; G.Y., M.P., P.C., and T.-Y.C. analyzed data; and G.Y., M.P., R.B., P.C., and T.-Y.C. wrote the paper. **Competing Interest Statement:** The authors have no competing interests to declare.

## Abstract

Vitamin B_12_ (cobalamin) is a high-value yet scarce cofactor required for various metabolic processes, making its efficient handling important for maintaining metabolic homeostasis. While the involvement of ATP:cob(I)alamin adenosyltransferases (MMAB) in the synthesis, delivery, and repair of 5’-deoxyadenosylcobalamin (AdoCbl) is well established, the kinetic mechanisms that regulate this process, particularly its negative cooperativity, remain poorly understood. Understanding these mechanisms is key to clarifying how MMAB efficiently uses AdoCbl, prevents resource wastage, and supports bacterial survival in nutrient-limited environments. Using single-molecule relative fluorescence (SRF) spectroscopy, we found that conformation-gated binding is the driving force behind MMAB’s preference for AdoCbl over hydroxocobalamin and is the underlying mechanism for negative cooperativity. This mechanism significantly slows down the binding of the second equivalent of AdoCbl, favoring the singly bound state. Our findings indicate that MMAB predominantly binds a single AdoCbl, optimizing the AdoCbl loading to methylmalonyl-CoA mutase. Additionally, our SRF approach also serves as a tool to explore other cofactor interactions, such as those between riboswitches and cobalamin derivatives, to provide insights into regulatory mechanisms of cobalamin sensing and gene regulation, which are crucial for bacterial adaptation to changing nutrient conditions.

**Significance Statement:** MMAB is important for B_12_-dependent propionate metabolism in bacteria. Our findings reveal that conformation-driven binding mechanism underlines the negative cooperativity of MMAB, as it favors the binding of the first AdoCbl while limiting further binding. The larger *k*_on_ for the first site, combined with similar unbinding rates for both sites, could provide a solution for optimizing cobalamin handling and minimize unnecessary waste. Our single-molecule fluorescence approach offers a powerful tool for investigating other dynamic cofactor interactions, providing new insights into regulatory mechanisms in bacterial metabolism.

## Introduction

Vitamin B_12_, or cobalamin, is a high-value cofactor for bacterial and mammalian physiology, and is trafficked by a network of chaperones that ensure its proper intracellular handling and targeting (1–4). Within this network, ATP:cob(I)alamin adenosyltransferase (MMAB or ATR) synthesizes 5’-deoxyadenosylcobalamin (AdoCbl) *in situ* (Fig. 1, *step* 1). In the absence of apo-methylmalonyl-CoA mutase (MMUT or MCM, *step* 2), AdoCbl and triphosphate (PPPi) can dissociate directly from MMAB, leading to potential wastage of a valuable cofactor. To minimize AdoCbl dissociation, MMAB initiates a homolytic reaction, forming cob(II)alamin that has a higher affinity for MMAB, and leading to cofactor sequestration instead. However, air oxidation of cob(II)alamin produces hydroxocobalamin (OHCbl), which then dissociates from MMAB (5–7). MMAB loads AdoCbl directly onto apo-MMUT to generate holo-MMUT (*step* 3), which catalyzes the conversion of methylmalonyl-CoA to succinyl-CoA (*step* 4), replenishing the TCA cycle and supporting energy metabolism (2, 3, 5, 6, 8–10).

**Figure 1.**
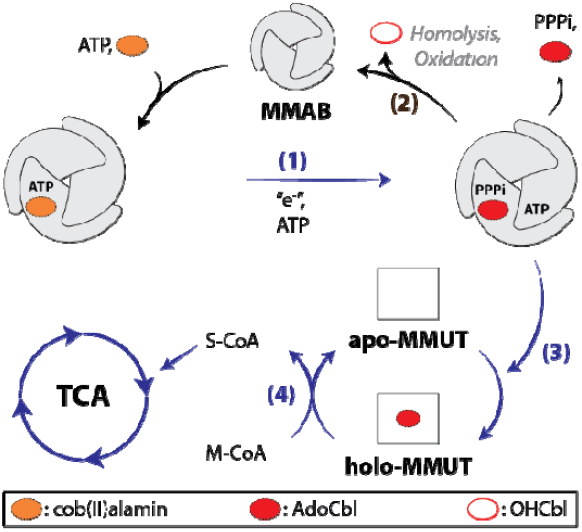
MMAB-catalyzed AdoCbl synthesis and transfer to MMUT. MMAB catalyzes AdoCbl synthesis from cob(II)alamin using ATP (step 1). Without apo-MMUT, AdoCbl undergoes homolysis and oxidation being converted to OHCbl, which dissociates from MMAB (step 2). Alternatively, MMAB transfers AdoCbl to apo-MMUT, forming holo-MMUT (step 3), which catalyzes the conversion of methylmalonyl-CoA to succinyl-CoA, replenishing the TCA cycle (step 4). MMAB can bind a maximum of two cob(II)alamin molecules. Consecutive binding and catalytic steps are omitted for clarity. This process is part of a larger protein network that optimizes B_12_ synthesis, trafficking, and utilization.

MMAB is a homotrimer and exhibits negative cooperativity, wherein binding of the first AdoCbl molecule decreases affinity for the second. The D180X truncation, corresponding to a clinical variant of human MMAB leads to loss of negative cooperativity and to inefficient cofactor transfer (11–14). Recent crystallographic studies have highlighted the importance of protein dynamics, particularly the role of mobile loops in in MMAB for mediating conformational changes during cobalamin binding and transfer (7). These findings suggest a potential link between structural flexibility and negative cooperativity, motivating elucidation of the kinetic basis of this regulatory mechanism. The detailed binding and unbinding rates that might explain how MMAB handles AdoCbl have not been described.

Investigating the interaction kinetics between MMAB and B_12_ derivatives at the ensemble level poses several challenges. The dynamic nature of these interactions can obscure transient states or rare conformations essential for enzyme function (15–17). Additionally, the homotrimeric architecture of MMAB complicates the measurement of cofactor stoichiometry. Traditional ensemble techniques like isothermal titration calorimetry, while informative, do not capture the full scope of these rapid and cooperative interactions (18, 19). To obtain a clearer, high-resolution view of kinetic behavior, more advanced methods such as single-molecule techniques (20–23) and/or advanced computational simulations (24–26) are required.

In this study, we have investigated the interaction kinetics of MMAB with B_12_ derivatives using a single-molecule relative fluorescence (SRF) approach. The instability of the native substrate, cob(II)alamin, which is highly prone to air oxidation (27, 28), complicates kinetic analysis and makes it difficult to directly study the substrate in its native form. To address this, we used a stable derivative of cobalamin, i.e., OHCbl, as a surrogate of the cob(II)alamin substrate, and the product, AdoCbl, to probe MMAB cofactor interactions. The SRF intensities enabled us to distinguish between MMAB states with 0-2 bound B_12_ equivalents, and to determine the transition times between these states. We find that a two-step kinetic model, derived from dwell-time distributions, effectively describes the interaction of MMAB with AdoCbl and OHCbl. Our kinetic model indicates that MMAB undergoes a conformational change before binding AdoCbl or OHCbl. The data reveals that the faster binding of the first B_12_ moiety, rather than faster unbinding of the second B_12_ moiety, is responsible for the observed negative cooperativity. The model further suggests that loss of negative cooperativity and impaired cofactor handling of the D180X MMAB variant could be due to a disruption of this binding mechanism. These findings offer a clearer understanding of the kinetic mechanisms governing AdoCbl binding to and release from MMAB.

## Results

### SRF trajectories visualize MMAB-AdoCbl interactions

The UV-visible absorption of AdoCbl makes it a strong fluorescence quencher (Fig. 2*A*), enabling the investigation of MMAB-AdoCbl interactions using fluorescence microscopy. To streamline the dye-labeling procedure and avoid complications, the cysteines in MMAB, which are not conserved, were converted to serine, yielding a Cys-less variant. In this background, Ser-99 and Ser-169 were subsequently mutated to cysteine for labeling (*SI Appendix*, section 1). We selected trimeric MMAB with a single equivalent of Alexa555 dye at Cys-99 to test the AdoCbl concentration-dependent fluorescence at the ensemble level (*SI Appendix*, section 2). The fluorescence intensity decreased with increasing AdoCbl concentration (Fig. 2*B*), indicating that AdoCbl binding could be probed by fluorescence quenching. To directly visualize MMAB-AdoCbl interaction events in real-time, we conducted Förster resonance energy transfer (FRET)-based SRF imaging (*SI Appendix*, section 3). We immobilized the Alex555-labeled MMAB on a glass slide, flowed AdoCbl solutions into the imaging chamber, and detected single-molecule fluorescence. The dwell times of the intensity fluctuations report on the MMAB-AdoCbl interaction kinetics.

**Figure 2.**
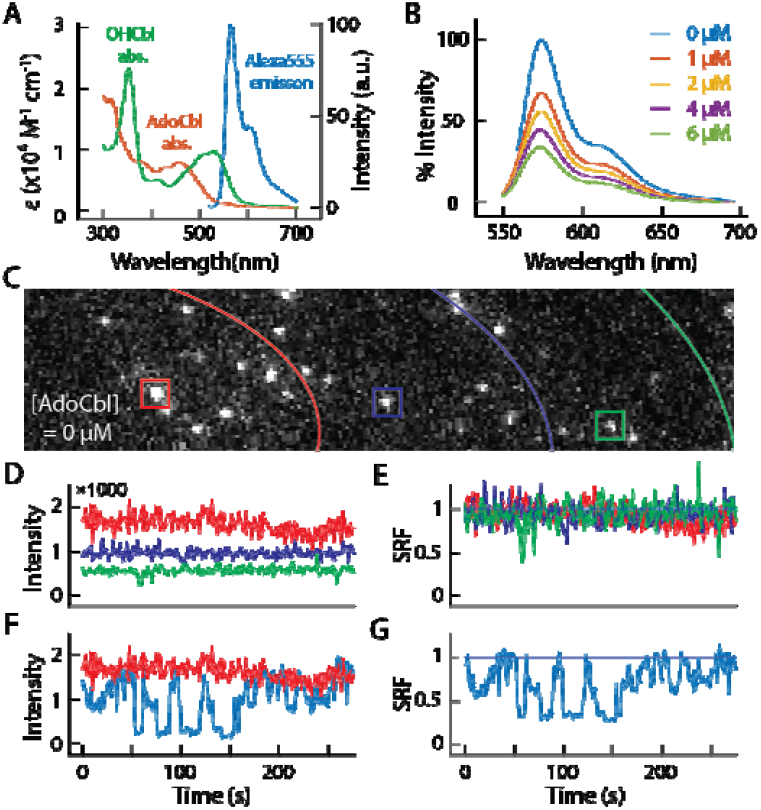
The generation of single-molecule relative fluorescence (SRF) trajectory. (*A*) Cobalamin absorption spectra overlayed with the Alexa555 emission spectrum. (*B*) Normalized fluorescence spectra of labeled MMAB in the solution with AdoCbl varying from 0 to 60 µM. (*C*) Fluorescence micrograph of Alexa555 labeled MMAB. The red, blue, and green curves mark the boundaries of the central, middle, and peripheral parts of the laser beam profile. (*D*) Intensity trajectories of red, blue, and green boxes in *C*. (*E*) Normalizing *D* by the average intensity of each trajectory gives the corresponding SRF trajectories. (*F*) Intensity trajectories of the red box in *C* at [AdoCbl] = 0 μM (red line) and 20 μM (blue line). (*G*) The normalized SRF trajectories of *F*.

To differentiate unbound (*MMAB*_U_), singly bound (*MMAB*_S_), and doubly bound (*MMAB*_D_) MMAB, we carefully removed the complications of inhomogeneous laser excitation and generated SRF trajectories (*SI Appendix*, section 4). Fig. 2*C* shows the single-molecule fluorescence image without AdoCbl. Each bright spot represents a single *MMAB*_U_ and maintains a relatively stable intensity over time (Fig. 2*D*). However, inhomogeneous laser excitation results in fluorescence signals with various intensities, even though all MMAB exist in an unbound state. To address this complication, we normalized each fluorescence signal of *MMAB*_U_ by its own average intensity to generate the SRF dataset (Fig. 2*E* - *G*). The SRF approach allows a direct combination and comparison of all fluorescence trajectories from different protein molecules. Since the quenching efficiency is directly related to the number of AdoCbl bound to MMAB, the SRF approach allows the differentiation of *MMAB*_U_, *MMAB*_S_, and *MMAB*_D_.

### SRF states reveal various MMAB-AdoCbl binding configurations

Fig. 3*A* shows SRF trajectories reporting interactions between MMAB and AdoCbl with different AdoCbl concentrations. The normalized SRF histograms (overall area equal to 1), generated from >200 SRF trajectories at each concentration, can be described by a combination of high (*F*_H_ ≈ 1.0), middle (*F*_M_ ≈ 0.77), and low (*F*_L_ ≈ 0.1) fluorescence states (Fig. 3*B*). We globally fit the histograms using a three-state Gaussian model. The center position and width of each Gaussian distribution were shared across all concentrations while allowing the relative populations of the three states to float. As AdoCbl increases from 0 to 20 µM, the *F*_H_ population decreased and shifted toward *F*_M_ while the *F*_L_ population grew continuously. A clear *F*_M_ peak appeared when AdoCbl was >40 µM, and the *F*_L_ peak gradually saturated. Thus, while *F*_H_ decreases, *F*_M_ and *F*_L_ increased and approached saturating populations (*SI Appendix section 5*, Fig. S4*C*).

**Figure 3.**
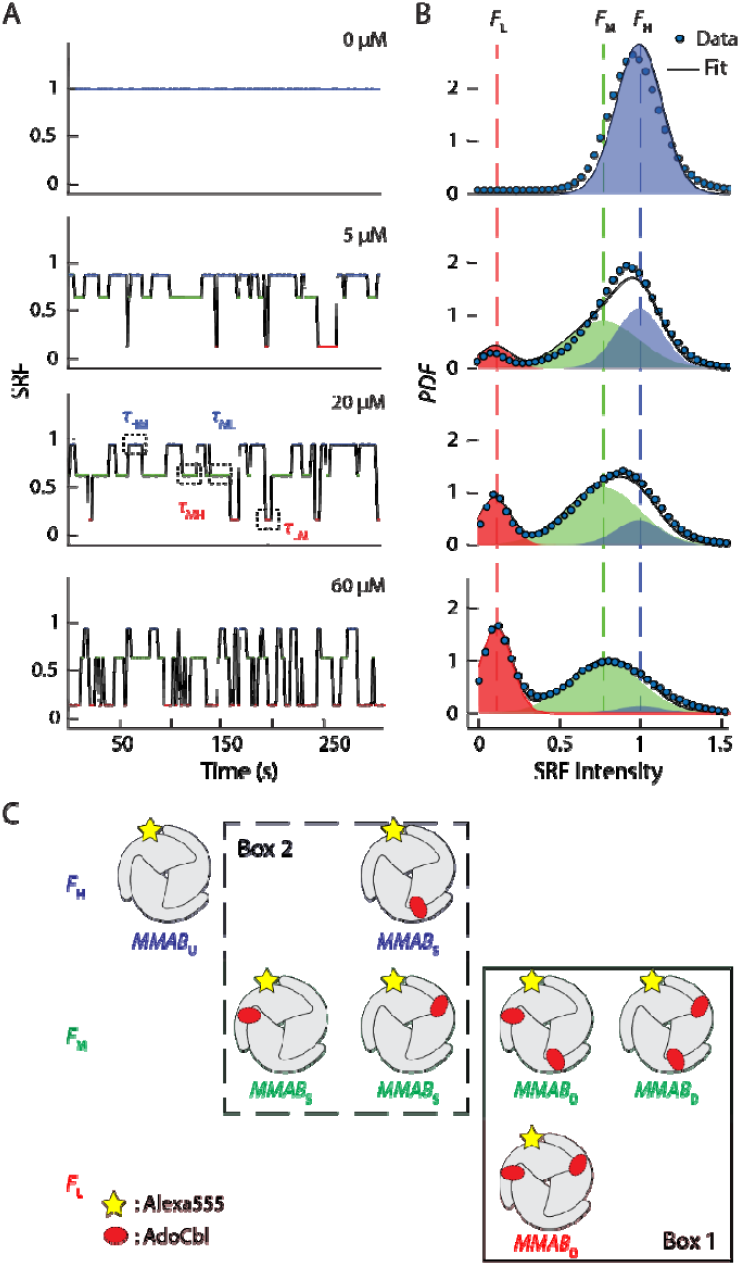
SRF states reveal various AdoCbl binding stoichiometries. (*A*) SRF trajectories at different AdoCbl concentrations. The solid lines are the corresponding transition trajectory generated from ebFRET. (*B*) SRF intensity histogram generated from *A* (blue solid circle) and the global fitting results (black line). The relative populations of *F*_H_, *F*_M_, and *F*_L_ are highlighted with blue, green, and red shades, respectively. (*C*) Assignment of various AdoCbl binding geometries of MMAB to the three SRF states. *MMAB*_U_, *MMAB*_S_, and *MMAB*_D_ represent MMAB under unbound, singly bound, and doubly bound state, respectively.

Using AdoCbl-dependent populations, we assigned SRF states to MMAB binding stoichiometries. Labeling MMAB with a single Alexa555 broke its trilateral symmetry and resulted in different binding configurations. Fig. 3*C* summarizes all possible MMAB binding configurations. In the absence of AdoCbl, the histogram shows a single state with *F*_H_ ≈ 1.0 (Fig. 3*B*, top), indicating that only *MMAB*_U_ contributed to the *F*_H_ state (Fig. 3*C*, top left). At AdoCbl = 60 µM, a saturating concentration, MMAB exists predominantly in the doubly bound form, *MMAB*_D_, giving three binding configurations. The ratio of *F*_M_ to *F*_L_ was ∼2:1 (Fig. 3*B*), indicating that two and one *MMAB*_D_ contributed to *F*_M_ and *F*_L_ states, respectively. Since the SRF intensity is dependent on the distance between the label and bound AdoCbl, we assigned the *MMAB*_D_ with two AdoCbl bound in the closest positions to the *F*_L_ state and the other two to the *F*_M_ state (Fig. 3*C*, Box 1). This assignment also indicates that the quenching efficiency of AdoCbl bound to the farthest site is insignificant. Application of this concept to the singly bound case led to the assignment of the *MMAB*_S_ species in the *F*_H_ and *F*_M_ states in a 1:2 ratio (Fig. 3*C*, Box 2). These assignments were also supported by another MMAB construct labeled at Cys-169 (*SI Appendix section* 5). From the above analysis, we associated *F*_H_ with *MMAB*_U_ + 1/3 *MMAB*_S_, *F*_M_ with 2/3 *MMAB*_s_ + 2/3 *MMAB*_D_, and *F*_L_ state with 1/3 *MMAB*_D_.

### Dwell-time analysis reveals an intermediate species and a kinetic model for MMAB-AdoCbl interactions

Most AdoCbl-induced transitions occur between *F*_H_ and *F*_M_ or *F*_M_ and *F*_L,_ while direct transitions between *F*_H_ and *F*_L_ states are occasionally observed (∼5%). This observation indicates that *MMAB*_D_ forms primarily through two sequential AdoCbl bindings and that simultaneous binding and unbinding of two AdoCbl is rare and thus not considered further. To quantify interaction kinetics, SRF trajectories were first analyzed using the empirical Bayes hidden Markov model analysis software, ebFRET, to generate transition trajectories (solid lines in Fig. 3*A, SI Appendix*, section 6) (29–31). We extracted transition dwell times from trajectories and generated the probability density functions (*PDF*) of dwell times (*SI Appendix*, section 7).

Τ_HM_ is the microscopic dwell time in *F*_H_ before transitioning to the *F*_M_ state. The overall average transition rate, ⟨Τ_HM_⟩^-1^, increased with increasing AdoCbl concentrations (Fig. 4*A*). However, the faster rate constant (i.e., steeper slope) in the low AdoCbl concentration region indicates a faster *MMAB*_U_ to *MMAB*_S_ than *MMAB*_S_ to *MMAB*_D_ transition. Interestingly, the distribution of Τ_HM_, instead of being a simple single-exponential decay, shows an exponential rise and decay, suggesting a kinetic intermediate in the *F*_H_ state (32, 33). This rise-and-decay feature is most visible at low (<10 μM) but vanishes at high AdoCbl concentrations (40 μM, Fig. 4*B*), suggesting that this intermediate is primarily associated with the *MMAB*_U_ to *MMAB*_S_ transitions. We posit this intermediate as a conformational change in *MMAB*_U_ instead of *MMAB*_S_ in the *F*_H_ state. Using a single-molecule interaction simulation (SMIS, *SI Appendix*, section 8) (24), we confirmed that MMAB undergoes a conformational change (i.e., the transition from *MMAB*_U1_ to *MMAB*_U2_) before the first AdoCbl binds.

**Figure 4.**
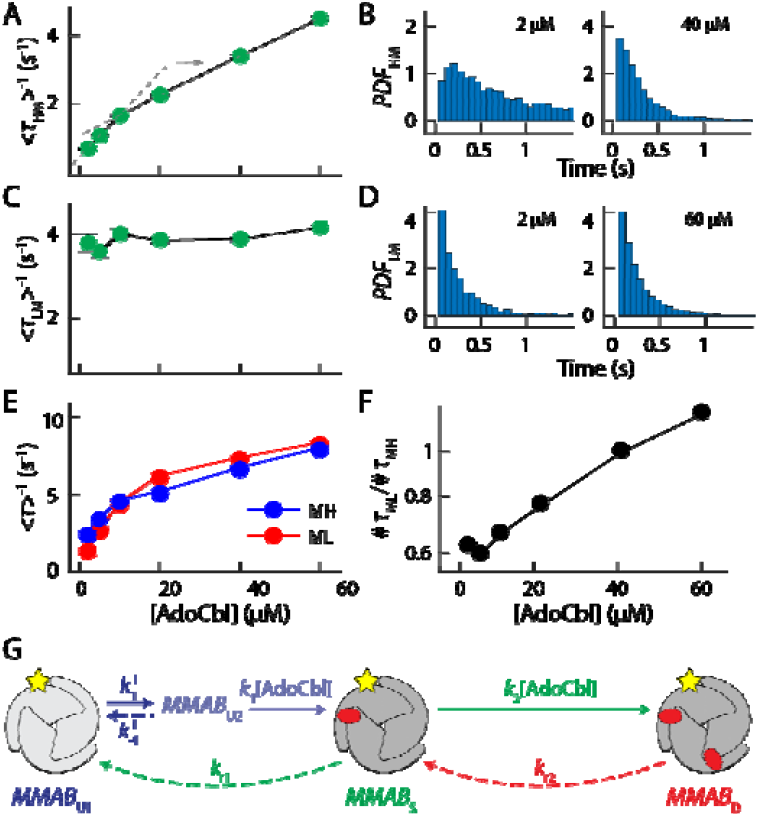
Microscopic dwell times inform a minimal kinetic model for MMAB-AdoCbl interactions. (*A*) AdoCbl dependent <Τ_HM_>^-1^ shows the averaged binding rate increases with AdoCbl concentrations. (*B*) *PDF*_HM_ under low and high AdoCbl concentrations. (*C*) AdoCbl-dependent <Τ_LM_>^-1^ shows that the unbinding rate from *MMAB*_D_ is AdoCbl independent. (*D*) Same as *B* but for *PDF*_LM_. (*E*) AdoCbl-dependent <Τ_MH_>^-1^ and <Τ_ML_>^-1^ show a similar dose dependence. (*F*) The ratio of transition events between Τ_ML_ and Τ_MH_ processes increases linearly with AdoCbl. (*G*) Proposed kinetic model describing MMAB-AdoCbl interactions. *MMAB*_U1_ needs to undergo a conformational change (i.e., transition to *MMAB*_U2_) before the first AdoCbl binding.

On the other hand, Τ_LM_ captures the transitions from the *F*_L_ to the *F*_M_ state and describes AdoCbl unbinding from the *MMAB*_D_. The ⟨Τ_LM_⟩^-1^ and *PDF*_LM_ remain the same across all AdoCbl concentrations (Fig. 4*C* and *D*). The dose-independent rates indicate that AdoCbl unbinds from *MMAB*_D_ via a typical unimolecular mechanism. Τ_ML_ and Τ_MH_, the microscopic dwell times of the *F*_M_ state before respectively transitioning to the *F*_L_ and *F*_H_ states, contain mixed AdoCbl binding and unbinding information for *MMAB*_S_ and *MMAB*_D_. Both ⟨Τ_ML_⟩^-1^ and ⟨Τ_MH_⟩^-1^ reveal the overall rates of all kinetic processes that start from the *F*_M_ state, which is supported by the fact that both ⟨Τ_ML_⟩^-1^ and ⟨Τ_MH_⟩^-1^ show identical AdoCbl dependence (Fig. 4*E*). The ratio of the number of transition events for Τ_ML_ and Τ_MH_ process (i.e., *#*_ML_/*#*_MH_) increases linearly with AdoCbl (Fig. 4*F*). Since the binding rate of AdoCbl to MMAB is expected to be scaled linearly with AdoCbl, the *#*_ML_/*#*_MH_ data indicate that AdoCbl unbinding from *MMAB*_S_ also occurs through a unimolecular mechanism.

The above kinetic analysis revealed that: (i) AdoCbl binds to MMAB mainly through a two-step binding model (reflected by rare direct transitions between *F*_H_ and *F*_L_); (ii) Binding kinetics of AdoCbl to MMAB show a clear AdoCbl dose dependence. The binding of the first AdoCbl equivalent requires the formation of a different conformational intermediate (reflected by the rise-and-decay feature of Τ_HM_ distribution), indicating that unbound MMAB equilibrates between at least two different conformations. This observation is also supported by the recent crystal structure showing that MMAB undergoes conformational changes upon binding AdoCbl or ATP (7). (iii) AdoCbl unbinding from both sites of MMAB follows an unimolecular mechanism (reflected by dose-independent ⟨Τ_LM_⟩^-1^ and dose-dependent *#*_ML_/*#*_MH_). Although it is clear that AdoCbl unbinding from *MMAB*_D_ leads to *MMAB*_S_, unbinding from *MMAB*_S_ can lead to either *MMAB*_U1_ or *MMAB*_U2_. We thus examined both kinetic models using SMIS and found that only the model in which AdoCbl unbinding leads to *MMAB*_U1_ describes all dwell time distributions (*SI Appendix*, section 9). Combining this information, we formulated a minimal kinetic model to describe the interactions between MMAB and AdoCbl (Fig. 4*G*).

### The kinetic model describes the complex kinetics of MMAB-AdoCbl interactions

Applying the MMAB-AdoCbl binding configurations (Fig. 3*C*) to a minimal kinetic model (Fig. 4*G*) resulted in a fairly complex system (Fig. 5*A*), precluding the feasibility of deriving an analytical solution for most dwell-time *PDF*s (i.e., *f*_HM_(Τ), *f*_ML_(Τ), and *f*_MH_(Τ)). *f*_LM_(Τ) is the only exception, because it only describes the unbinding step for *MMAB*_D_ 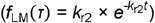. Fitting the distributions of Τ_LM_ with *f*_LM_(Τ) crossing all AdoCbl concentrations revealed the unbinding rate constants *k*_r2_ (Fig. 5*B*). The derivation of either *f*_HM_(Τ), *f*_MH_(Τ), or *f*_ML_(Τ) was too complicated due to the ill-defined ratio of the initial species (i.e., *F*_H_ and *F*_M_ states). To circumvent this challenge, we defined and extracted two additional dwell times, Τ_LMH_ and Τ_LML_. Τ_LMH_ and Τ_LML_ represent the subsets of Τ_MH_ and Τ_ML_, respectively, which include only the dwell time starting from the *F*_L_ state. This selection results in Τ_LMH_ and Τ_LML_ having a well-defined initial state (i.e., *MMAB*_S_ in the *F*_M_ state, Fig. 5*A*) and enables derivations of *f*_LML_(Τ) and *f*_LMH_(Τ) (*SI Appendix*, section 10). The global fitting of *PDF*_LML_ and *PDF*_LMH_ with *f*_LML_(Τ) and *f*_LMH_(Τ) (Fig.5*C*) yielded *k*_2_ and *k*_r1_. To quantify 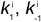 and *k*_1_, we used SMIS to test a large range of 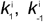 and *k*_1_ and obtained a solution set that satisfactorily described our data (Fig. 5*D*) (24). Based on the kinetic model, we derived the unbinding equilibrium constants *K*_D1_ and *K*_D2_ (*SI Appendix*, section 11).

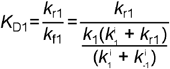

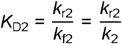

**Figure 5.**
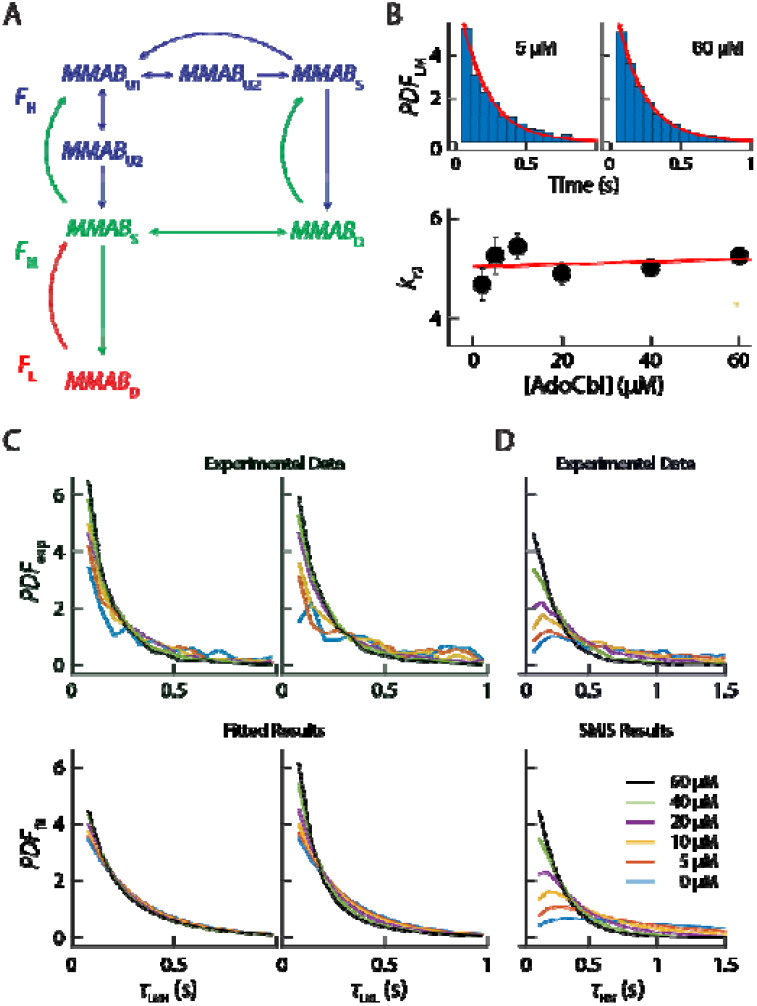
Kinetic parameters for MMAB-AdoCbl interactions. (*A*) The minimal kinetic model associates different MMAB species to SRF states. (*B*) Fitting the *PDF*_LM_ (blue bar) by a single-exponential function (red curve) reports *k*_r2_. The AdoCbl independent fitted *k*_r2_ suggests AdoCbl unbinds from *MMAB*_D_ through single-step dissociation (*C*) The global fitting of *PDF*_LML_ and *PDF*_LMH_ with *f*_LML_(Τ) and *f*_LMH_(Τ) reports *k*_2_ and *k*_r1_. (*D*) The extraction of, _-_, and *k*_1_ was achieved using SMIS. The simulated Τ_HM_ distributions (bottom) satisfactorily describes the experimental data (top).

*k*_f1_ and *k*_r1_ represent the forward binding and reverse unbinding rate constants for transitions between *MMAB*_U_ and *MMAB*_S,_ while *k*_f2_ and *k*_r2_ for transitions between *MMAB*_S_ and *MMAB*_D_. Table 1 summarizes the extracted rate constants and reports values for *K*_D1_ = 5.6 ± 2.9 μM and *K*_D2_ = 41.7 ± 13.9 μM. *K*_D2_ is 7-fold larger than *K*_D1_, which is significantly larger than the 3.3-fold difference predicted for a system exhibiting no cooperativity (*SI Appendix*, section 12), indicating the negative cooperativity.

**Table 1.**
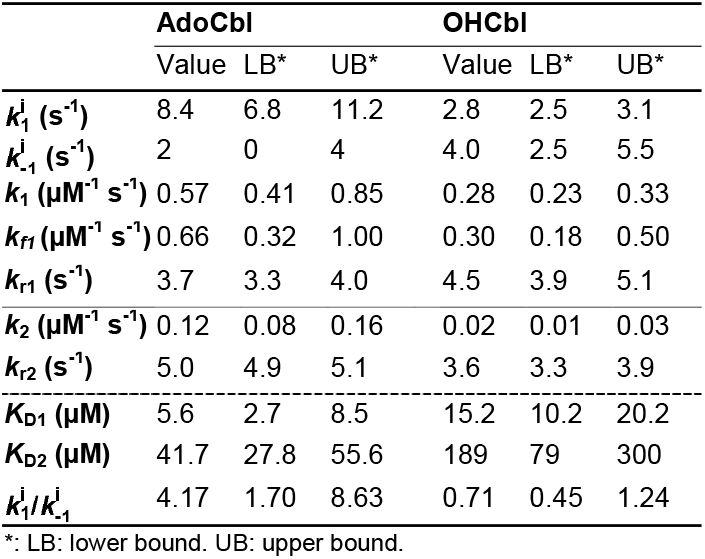
Kinetic parameters for MMAB-B_12_ interactions.

### HOCbl interacts with MMAB using a kinetic model similar to AdoCbl

HOCbl binding to MMAB showed a stronger quenching than AdoCbl because of its larger FRET spectral overlap with the Alexa dye (Fig. 2*A*). SRF intensity histograms identified three fluorescence states (Fig. 6*A*), which were used to determine the HOCbl-dependent populations (Fig. 6*B*) and assign possible binding configurations (Fig. 6*C* and *SI Appendix*, section 13). By carefully analyzing the dwell time distributions (Fig. 6*D*), we proposed a similar minimal kinetic model to describe the MMAB-HOCbl interactions (Fig. 6*E* and *SI Appendix*, section 14). MMAB also required a conformational change (*MMAB*_U1_ to *MMAB*_U3_) to bind the first equivalent of HOCbl (Fig. 6*E*). We derived analytical equations for all dwell times (i.e., Τ_HM_, Τ_MH_, Τ_ML,_ and Τ_LM_, *SI Appendix*, section 15), applied equations to fit the dwell-time distributions, and summarized the rate constants in Table 1. Notably, the conformational change rate constants 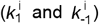 of MMAB_U1_ for HOCbl are very different from those of AdoCbl. These differences suggest that *MMAB*_U1_ adopts a different conformation, which was designated *MMAB*_U3_. The binding affinity of MMAB to HOCbl is much weaker than that to AdoCbl (*K*_D1_ = 15.2 ± 5.0 μM and *K*_D2_ = 189 ± 110 μM), but they still show a negative cooperativity originated from the slower second binding.

**Figure 6.**
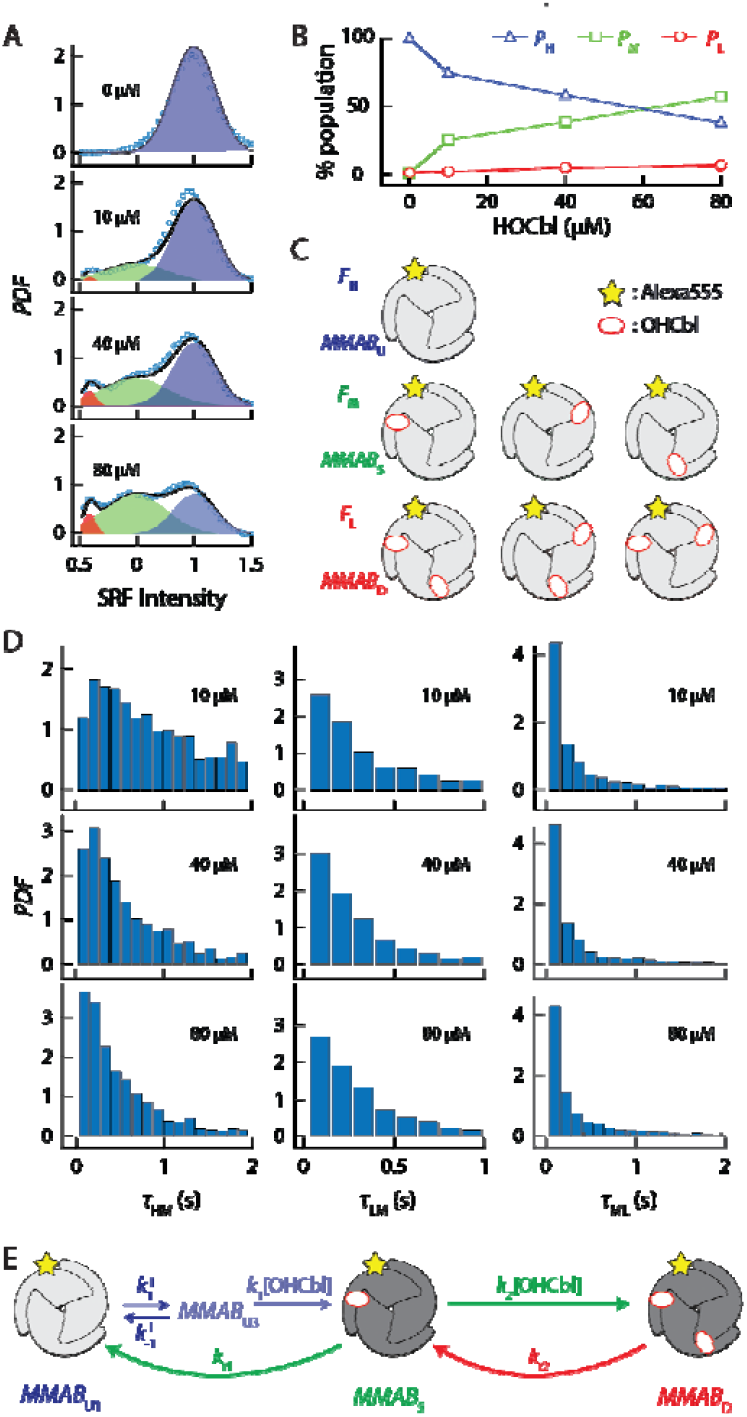
Kinetic analysis of MMAB-OHCbl interactions. (*A*) OHCbl-dependent SRF intensity histogram. (*B*) OHCbl-dependent populations of *F*_H_, *F*_M_, and *F*_L_ states. (*C*) Assignment of all possible OHCbl binding configurations of MMAB (*MMAB*_U_, *MMAB*_S_, and *MMAB*_D_) to the three SRF states. (*D*) Experimental dwell time distributions of Τ_HM_, Τ_LM_, and Τ_ML_. (*E*) Proposed kinetic model describing MMAB-OHCbl interactions. *MMAB*_U1_, *MMAB*_S_, and *MMAB*_D_ represent MMAB in the unbound, singly and doubly bound states, respectively. *MMAB*_U1_ needs to undergo a conformational transition to *MMAB*_U3_ before binding the first equivalent of OHCbl.

## Discussion

### Kinetic model describes ensemble results

Using the SRF assay, we described a two-step kinetic model and quantified the kinetic and thermodynamic constants for interactions between MMAB and cobalamin. Before MMAB binds the first equivalent of cobalamin, it must adopt a specific conformation (*MMAB*_U2_ and *MMAB*_U3_). Binding of the first cobalamin moiety slows down binding of the second, leading to negative cooperativity. Unbinding of cobalamin from MMAB follows a unimolecular pathway and is independent of cobalamin concentration. Using the extracted rate constants (Table 1) and kinetic model (Fig. 4*G* and Fig. 6*E*), we simulated the stopped-flow differential absorption traces for AdoCbl binding to apo-MMAB (*SI Appendix*, section 15), which captured the AdoCbl dependence of differential ensemble absorption spectra reported previously,(6) thus validating our model.

### D180X loses negative cooperativity due to reduced *k*_on_ for the first substrate

In wild-type MMAB, negative cooperativity could stem from either a larger *k*_on_ for the first substrate relative to the second, or a larger *k*_off_ for the second substrate. Our kinetic data ruled out the second possibility, as the unbinding rates were similar across the binding sites. Therefore, the D180X mutation likely disrupted the first substrate binding event by reducing *k*_on_. This slower initial substrate capture is reflected in a 2-fold higher *K*_M_, as the enzyme now requires more substrate to reach half-maximal activity. The threefold lower *k*_cat_ is a direct consequence of inefficient turnover, due to inefficient binding of the first substrate equivalent, which slows down the overall catalytic cycle. The decrease in *k*_on_ corresponded to an increase in *K*_d_, explaining the observed 400-fold reduction in AdoCbl affinity.

### Biological implications of the kinetic model

Since the intracellular concentration of B_12_ derivatives is relatively low (nM to low µM), efficient substrate capture is important for the catalytic efficiency of ATR. A larger *k*_on_ for binding at the first site allows ATR to rapidly bind cob(II)alamin and convert it to AdoCbl, facilitating cofactor synthesis under substrate-limited conditions. Previous studies have shown that when MMAB is doubly loaded, ATP selectively ejects AdoCbl from the weaker-binding second site while retaining the tightly bound AdoCbl at the first site (34). This selective ejection minimizes premature cofactor loss into solution and results in MMAB retaining one AdoCbl equivalent, potentially serving as a functional reserve. Negative cooperativity slows down the binding of a second equivalent of AdoCbl, which contributes to disfavoring back transfer of the cofactor from MMUT to MMAB. If both sites were to bind cob(II)alamin and convert it to AdoCbl at similar rates, the balance between cofactor synthesis and transfer could be disrupted, increasing the likelihood of premature AdoCbl release into solution. Thus, the combination of negative cooperativity and ATP-driven AdoCbl transfer appears to represent a solution for balancing cob(II)alamin availability with the AdoCbl demand.

### Broader applicability of SRF

Our SRF approach revealed the dynamic interactions and binding kinetics of MMAB with B_12_ derivatives and provided insights into the basic mechanism of cofactor handling by MMAB. The SRF approach holds the potential for broader applications, for example, to study the interaction between riboswitches and cobalamin derivatives (35–37), offering a powerful tool for probing the regulatory mechanisms governing cobalamin sensing and gene regulation in bacterial systems.

### Materials and Methods

Materials and methods are described in *SI Appendix*, section 1 to section 16. These include Alexa-labeled MMAB and functional validation, experimental setups for single-molecule and ensemble fluorescence imaging, procedures for image and data analyses, experimental dwell-time distribution generation, dwell-time analytical solution derivation based on kinetic models, and stopped-flow differential absorption simulations.

## Acknowledgments

The authors thank the Biotechnology Resource Center for protein mass spectrometry analysis. The authors gratefully acknowledge financial support from NIH (R35GM133505 (T.-Y.C.), R01GM109993 (P.C.), and R01DK45776 (R.B.)) and the University of Houston.

