## Supporting Information for "Conformation-gated binding drives negative cooperativity in ATP:cob(I)alamin Adenosyltransferase for optimized cobalamin handling"

**This PDF file includes:**

Supporting text

Figures S1 to S12

Tables S1 to S3

SI References

**Table of content**

1. **Preparation of Alexa555-labeled MMAB** **3**
2. **Validation of fluorescence quenching assay at the ensemble level** **5**
3. **Single-molecule fluorescence microscope setup** **6**
4. **Generation of single-molecule relative fluorescence (SRF) trajectories** **7**
5. **Validation of the SRF state assignment** **9**
6. **Extraction of dwell-times from SRF trajectories using ebFRET** **11**
7. **Generation of probability density functions of dwell times** **12**
8. **$MMAB_s$ is not the intermediate in the $F_H$ state** **13**
9. **$MMAB_s$ must return to $MMAB_{U1}$ when AdoCbl dissociation occurs** **15**
10. **Derivations of $f_{LMH}(\tau)$ and $f_{LML}(\tau)$** **16**
11. **Derivation of $K_{D1}$ and $K_{D2}$** **19**
12. **SRF state assignments and subpopulation analysis for MMAB-OHCbl interactions** **21**
13. **Minimal kinetic model for MMAB-OHCbl interactions** **23**
14. **Extraction of kinetic rate constants for MMAB-OHCbl interactions** **25**
15. **Simulation of stopped-flow differential absorption** **27**
16. **Reference** **29**

### 1. Preparation of Alexa555-labeled MMAB

To visualize MMAB through a fluorescence microscope, we used mutagenesis to introduce a unique cysteine for fluorescent probe labeling. The *Methylobacterium extorquens* MMAB contains two natural non-conserved cysteines (C64 and C154) in each subunit. Both cysteines are less surface-exposed and, thus, not ideal for serving as labeling sites. To avoid possible complications in the dye-labeling procedure, we mutated these two cysteines to serine to create a Cys-less MMAB. We then identified S99 and S169 in two-loop regions to introduce cysteine for labeling (Fig. S1*A*). These two sites are not-conserved, solvent-exposed, do not participate in H-bonding to stabilize the protein structure, and are distant from the AdoCbl binding sites.

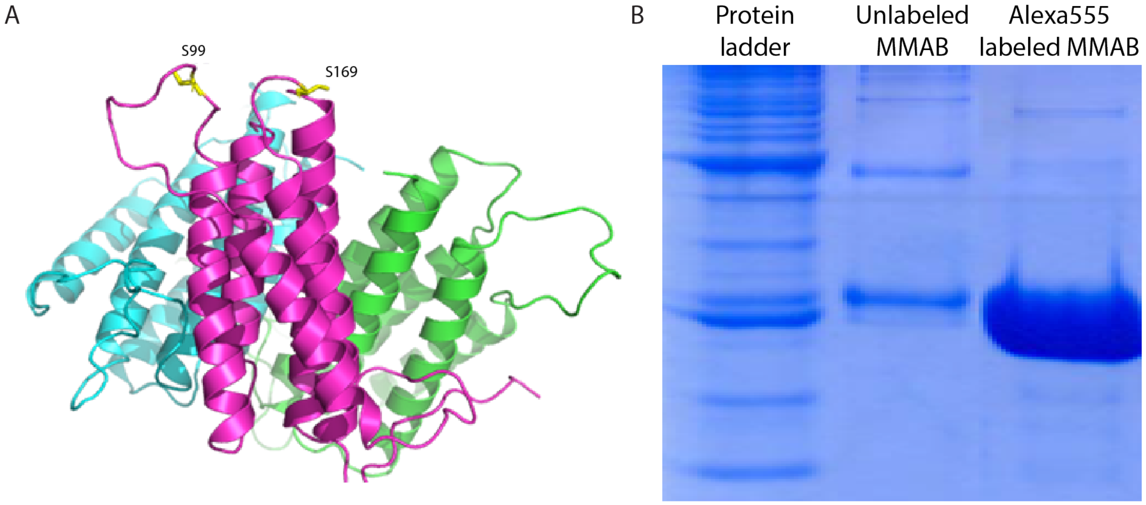


**Supplementary Fig. S1. MMAB crystal structure and characterization.** (*A*)Two sites for Alexa555-labeleing of MMAB. S99 is in the loop between the α2 and α3 helices. S169 is in the loop between the α4 and α5 helices. (*B*) SDS-PAGE gel of S99 construct with Coomassie blue staining after MonoQ column purification. Column from left to right is the protein ladder, unlabeled MMAB, and Alexa555-labeled MMAB, respectively.

Both MMAB variants were active in catalyzing AdoCbl synthesis and showed no precipitation at room temperature for >2 h (Table S1). S99C MMAB maintained 94% catalytic activity and, like wild-type MMAB, released only one equivalent of AdoCbl into solution. In contrast, S169C MMAB exhibited 41% of wild-type activity while also releasing only one equivalent of AdoCbl. For the remainder of the study, we focused on S99C MMAB.

**Table S1. Activity results of the wild-type MMAB and two MMAB constructs.**

| | **WT MMAB** | **Cys-less S99C** | **Cys-less S169C** |
|---|---|---|---|
| **$k_{cat}$ (min$^{-1}$)** | 0.87 | 0.82 | 0.36 |
| **Fold change** | | 0.94 | 0.41 |
| **Max % AdoCbl Release** | 53 | 44 | 46 |
| **Stability** | Stable at RT | Stable at RT >2h (90 µM) | Stable at RT >2h (100 µM) |

The MMAB mutants were cloned in a pET28 vector and overexpressed in *E. coli* BL21 (DE3), growing in 20 g/L LB medium, containing 30 mg/L kanamycin and 34 mg/L chloramphenicol or 0.05 mg/mL kanamycin at 37 ˚C. After $OD_{600nm}$ reached 0.6, 1 mM IPTG was added to the solution to induce expression for 4 h. Cells were then centrifuged and resuspended in Buffer 1 (50 mM sodium phosphate buffer, pH 8.0, containing 0.3 M KCl, 20 mM imidazole, 0.1 mM PMSF, 5% glycerol), then 100 mg lysozyme was added and stirring was continued at 4 ˚C for 30 min. Cells were sonicated, and the supernatant was collected following centrifugation to remove cell debris. The final supernatant was passed through a 0.2 µm syringe filter and purified by HPLC using a His-trap column as described previously (1).

The MMAB variants were labeled with Alexa555 using maleimide chemistry. Before labeling, MMAB was reduced by mixing with 5 µL TCEP (200 mM) and 200 µL MMAB (0.45 mM) in Buffer 1 to achieve final concentrations of 4.88 mM and 0.44 mM, respectively. The solution was incubated at 4 ˚C overnight. The Alexa555-Maleimide/Bio-PEG-Maleimide solution was prepared by mixing 13.3 µL Alexa555 in DMSO (30 mM) with 3.2 µL Bio-PEG-Maleimide in DMSO (250 mM). Reduced MMAB (205 µL, 0.44 mM) was mixed with 4 µL of Alexa555-Maleimide/Bio-PEG-Maleimide mixture in a final volume of 209 µL. The mixture was flushed with $N_2$ for 1 min to exclude $O_2$, and incubated at room temperature for 1.5 h or overnight at 4 ˚C. The labeling reaction was performed in the dark to protect the fluorescent dye Alexa555. Labeled MMAB was passed through a Superdex peptide column (GE Healthcare, Superdex Peptide 10/300 GL) and MonoQ anion column (GE Healthcare, Tricorn Mono Q 5/50 GL) with elution buffer (50 mM sodium phosphate buffer, pH 8.0, containing 0.3 M KCl, 150 mM imidazole, 0.1 mM PMSF, 5% glycerol) to obtain Alexa55dye-labeled MMAB, which was confirmed by SDS-PAGE gel (Fig. S1*B*).

### 2. Validation of fluorescence quenching assay at the ensemble level

To validate that the interactions between dye-labeled MMAB and cobalamin indeed leads to a fluorescence intensity decrease, we quantified the fluorescence intensity of MMAB solutions with different cobalamin concentrations. A control experiment (solution with dye instead of dye-labeled MMAB) was performed in parallel to ensure the quenching was indeed due to MMAB-cobalamin interaction rather than random collisions between the dye and cobalamin.

We labeled the S99C MMAB with a FRET donor dye, Alexa555 or Cy3, and used AdoCbl as the acceptor. Note that Alexa555 and Cy3 have very similar emission spectra, with the emission maximum centered around 555 nm (Fig. S2*A*). The rationale for using AdoCbl instead of OHCbl is that AdoCbl has a smaller spectral overlap (i.e., lower quenching efficiency, Fig. 1*A*). We reasoned that if AdoCbl shows significant quenching, this approach was likely to be even more effective for probing MMAB-OHCbl interactions.

Fig. S2*B* shows the fluorescent spectrum of 0.5 µM Cy3-labeled S99C MMAB mixed with AdoCbl, (0 to 6 µM). Due to the interaction of the donor MMAB-Cy3 and the acceptor AdoCbl, the fluorescent signal decreases with increasing AdoCbl concentrations and reaches ~30% of the initial intensity at [AdoCbl] = 6 µM. In contrast, the Cy3 control experiment shows the fluorescent spectrum of 0.5 µM Cy3 mixed still maintains ~ 80% of the initial intensity. The result from the control experiment indicates that random collisions between Cy3 and AdoCbl decrease the fluorescent signal only slightly. These results indicate that the fluorescent quenching assay can detect the interaction of MMAB and AdoCbl. We thus applied this concept and investigated the MMAB-cobalamin interactions via a single-molecule fluorescence quenching assay.

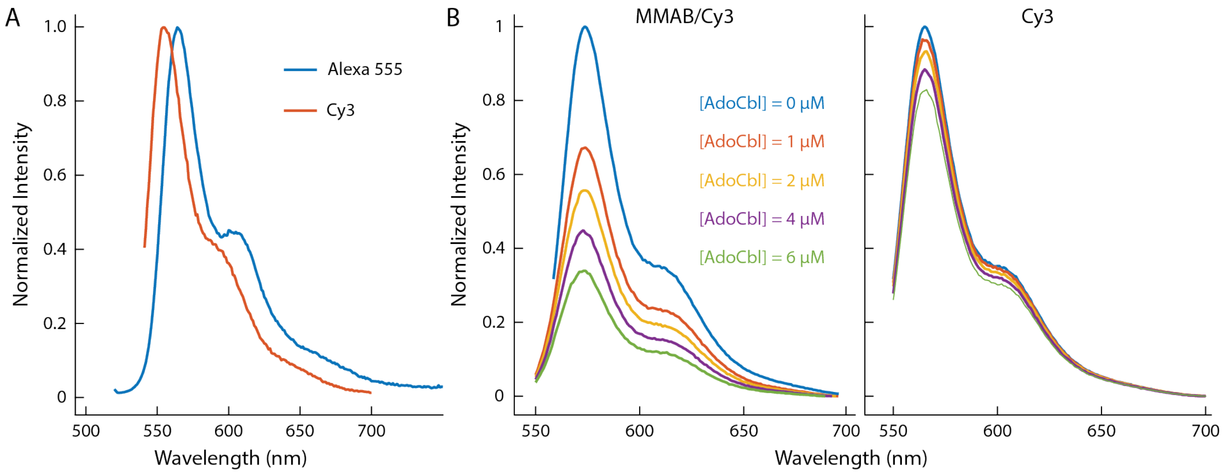


**Supplementary Fig. S2**. **Ensemble fluorescence quenching assay.** (*A*) The emission spectrum of Alexa555 and Cy3. (*B*) Normalized intensity of 0.5 µM MMAB-Cy3 (left) and Cy3 (right) in the solution with [AdoCbl] varying from 0 to 6 µM.

### 3. Single-molecule fluorescence microscope setup

Single-molecule relative fluorescence (SRF) experiments were performed on an Olympus IX71 inverted microscope using a prism-type total internal reflection fluorescence (TIRF) configuration. In the excitation path, the 532 nm laser (CrystaLaser) passed a quarter waveplate (Thorlabs AQWP05M-600) to ensure circularly polarized excitation. The laser excited labeled MMAB under TIRF geometry at a power of ~6-7 mW across a 68 × 137 $\mu m^2$ area. The fluorescent signal from Alexa555 was collected through a 60×, NA = 1.2, water-immersion objective (Olympus) and filtered by an HQ550LP filter (Chroma) before entering the EMCCD camera (Andor iXon). The image of Alexa555 fluorescence intensity was obtained using Andor iQ software. The movie was acquired with an image integration time of 32 ms, and the typical movie length is ~1-2 min. A homemade flow chamber was prepared by fixing a clean coverslip on the clean quartz slide as reported (2). The surface of the quartz slide was functionalized with biotinylated albumin (Sigma) and neutravidin. The sample containing the labeled MMAB was incubated at ~30 pM for 5–15 min to ensure proper sample density for SRF measurements. AdoCbl solutions with various concentrations (0-60 µM) flow into the flow chamber with a flow velocity of 10 µL/min. A photoprotection system (1 mM Trolox and 10 mM cysteamine) was added into the AdoCbl solution right before each experiment to prevent the photoblinking and photobleaching of the fluorescent probe.

### 4. Generation of single-molecule relative fluorescence (SRF) trajectories

After identifying the single labeled MMAB spots from the fluorescence micrograph, we generated the intensity trajectory by integrating the pixel intensity of a 3 × 3-pixel area from the center of each spot (Fig. S3*A*). Only trajectories that showed a single step photobleaching event were analyzed. Due to the Gaussian profile of the excitation laser (Fig. S3*B*), spots in the center area show higher intensities than those in the peripheral regions. This intensity heterogeneity prevents direct intensity comparison and limits fluorescence quenching assay usage. To address this issue, we further normalized each intensity trajectory and generated the SRF trajectory for subsequent analysis using home-built Matlab functions following the steps below:

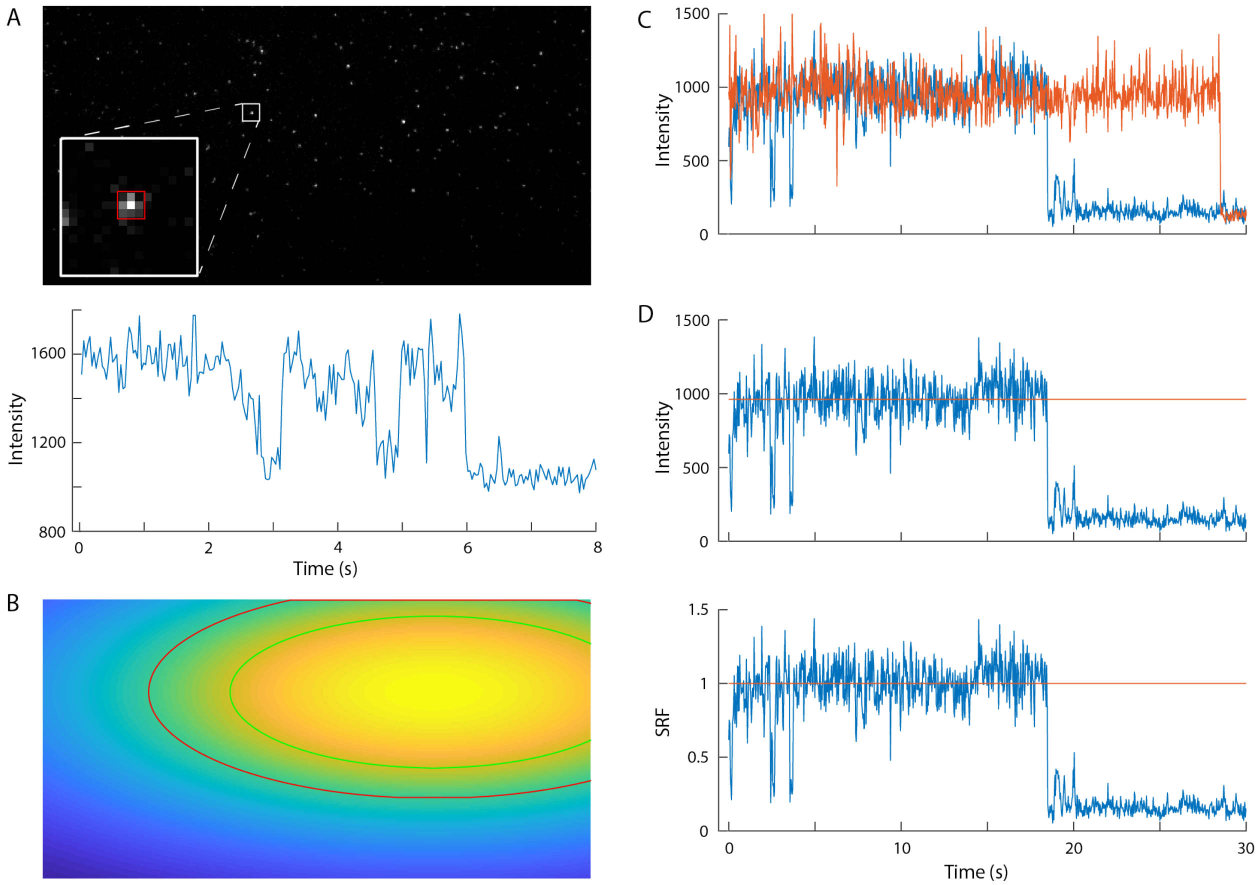


**Supplementary Fig. S3**. **Generations of SRF trajectory.** (*A*) Generation of intensity trajectory from the single-molecule micrograph. The pixel intensity of a 3 × 3-pixel area from the center of each spot (red square of the top) was integrated over time to generate the intensity trajectory (bottom). (*B*) Laser beam profile under TIRF-illumination geometry. (*C*) Target (blue) and reference (green) trajectory were used to generate the SRF trajectory. (*D*) Conversion from the intensity trajectory (top) to the corresponding SRF trajectory (bottom) is done by normalizing the intensity trajectory by the average intensity of the reference (top, red curve).

(i) We first conducted single-molecule fluorescence microscopy to collect dye-labeled MMAB intensities under cobalamin-free conditions. Without the complication of quencher (i.e., cobalamin), all singly labeled MMAB are expected to have identical intensities. When examining the intensity distribution of these singly labeled MMAB, we observed an excellent agreement with the laser excitation profile. We thus normalized the intensity trajectories by its mean intensity using the data without the photobleaching part and generated the SRF trajectories.

(ii) To generate the SRF trajectories for [AdoCbl] varying from 2-60 μM, we normalize their intensity trajectories with the selected intensity trajectory under AdoCbl-free conditions. We grouped the spot based on its distance to the center of the beam profile. We considered these spots under similar excitation power density and thus similar single-molecule intensity. For each target spot, we searched for reference spots in the same group under AdoCbl-free condition for normalization (Fig. S3*C*). We generated the final SRF trajectory by dividing the target trajectory by the mean intensity of the reference spot (Fig. S3*D*).

### 5. Validation of the SRF state assignment

To validate the assignment of SRF states, we further analyzed the SRF distribution of another construct, MMAB S169C (Fig. S4*A*).

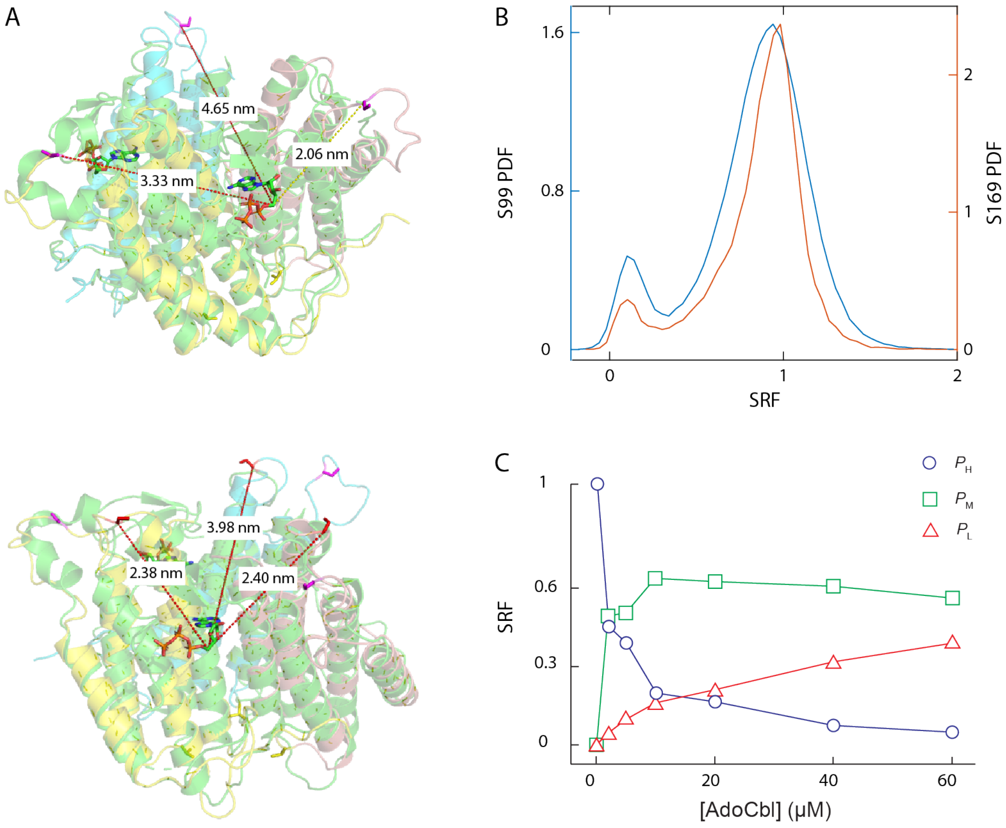


**Supplementary Fig. S4. Distances between dye and cobalamin binding sites, SRF distributions, and populations of [AdoCbl] dependent SRF states.** (*A*) Distances measured from the two labeling sites, S99 (left) and S169 (right), to three ATP binding sites. *M. extorquens MMAB structure* was aligned with human MMAB (green, pdb code: 2IDX), in which the ATP binding sites are resolved. Since the AdoCbl and ATP binding sites are in close proximity, distance between labeling site (S99 or S169) to ATP binding site are used to estimate the quenching efficiency and explain the experimental SRF distributions. (*B*) SRF distributions of S99C and S169C constructs under [AdoCbl] = 10 μM. Both shows similar distribution, indicating the farthest site did not contribute the fluorescence intensity decrease. The similar distances between the labeling site and two ATP binding sites, S169C construct shows a narrower SRF distributions compared to the S99C. (*C*) the relationship between SRF population and [AdoCbl], which was obtained through global fitting using a three-gaussian model with shared peak locations and width. As [AdoCbl] increases, the population of the high state ($P_H$) decreases, while the middle ($P_M$) and low ($P_L$) states increase and eventually reaches saturation.

Since quenching through FRET is distance sensitive, one can put the fluorescent tag at different locations and predict how the FRET states change. Table S2 summarized the distances between labeling sites (site 99 and 169) and the ATP binding site, a site that is in close proximity to AdoCbl binding sites. Both labeling sites are far away from one of the AdoCbl binding sites (site 3). Since the distances (4.65 nm for S169C and 3.98 nm for S99C) are much larger than the expected $r_0$ (2.0 nm), we expect the AdoCbl bound at this site would only slightly quench the fluorescence and would not be distinguishable from the unquenched Alexa555. However, the other two AdoCbl binding sites (site 1 and site 2) are closer to the labeling sites and expected to lead to significant fluorescence quenching. For the S99C labeling site, the distance to site 1 and site 2 is 2.06 nm and 3.33 nm, respectively.

**Table S1. Estimated distances between labeling and ATP sites**

| Distance to | Site 1 (nm) | Site 2 (nm) | Site 3 (nm) |
|---|---|---|---|
| S99 | 2.1 | 3.3 | 4.65 |
| S169 | 2.4 | 2.4 | 4 |

At [AdoCbl] = 10 µM condition, the SRF histogram shows a very broad peak centered ~ 0.92. Based on our assignment, this broad distribution mainly originated from a mixture of two doubly bound and three singly bound MMAB. For the S169C labeling site, the distance to site 1 (2.40 nm) is almost the same as that to site 2 (2.38 nm). If our assignment is correct, one would expect the two doubly bound and two singly bound *MMAB*s to show similar quenching efficiency at [AdoCbl] = 10 µM condition. Fig. S4*B* shows the SRF histogram for both constructs at [AdoCbl] = 10 µM. The S169C construct shows a much narrower peak than the S99C one, supporting our SRF state assignment.

The subpopulation analysis of the MMAB S99C mutant also supports this assignment. At [AdoCbl] = 2 µM (Fig. S4*C*), MMAB dominantly exists either in the $MMAB_U$ or $MMAB_S$. Since the $MMAB_U$ only contributes to the $I_H$ state, the population ratio between the middle ($P_M$) and low ($P_L$) states dictates the contribution of $MMAB_S$. The fact that $P_M/P_L$ being larger than three indicates no single-bound species contributes to the $I_L$ state. The trace $P_L$ informs the $MMAB_D$ is ~ 4%; therefore, $MMAB_D$ contributes 8% to $I_M$ population. This means ~ 42% of the $P_M$ (i.e., 50 − 8% = 42%) must be from the $MMAB_S$. The $P_M$ is a comparable amount to the $P_H$ (i.e., 46%), which is only possible when three $MMAB_S$ contribute to $I_H$ and $I_M$ states with a ratio of 1/2.

### 6. Extraction of dwell-times from SRF trajectories using ebFRET

Microscopic dwell times of each SRF trajectories were obtained using the ebFRET software (http://ebfret.github.io/) with the number of fluorescent states set to three (3). The ebFRET is a software that use empirical Bayes hidden Markov model analysis to analyze noisy trajectories and extract the microscopic dwell times.

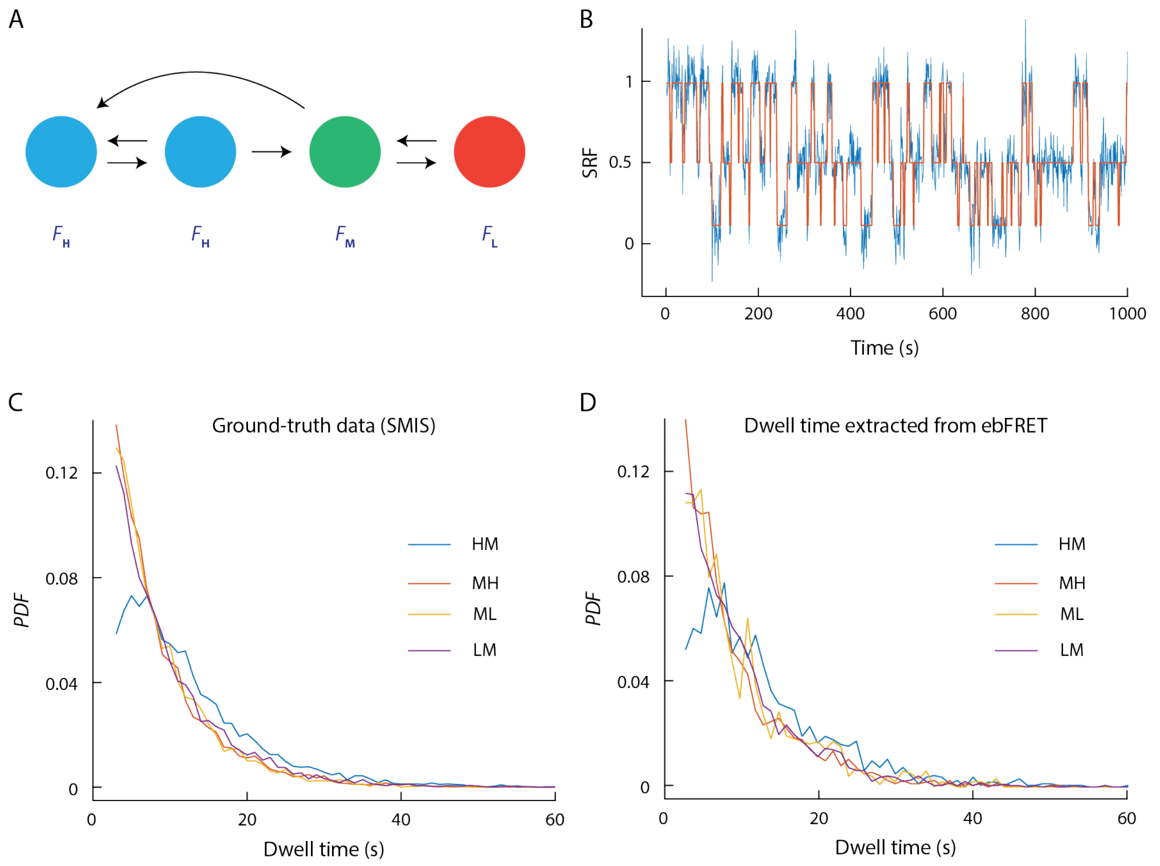


**Supplementary Fig. S5. ebFRET reasonably extract the dwell time distributions.** (*A*) Kinetic model for generating simulated single molecule trajectories. The two blue, green, and red circle represent the fluorescent high, middle, and low states, respectively. (*B*) The overlay of the simulated noisy ground true (blue) and the ebFRET extracted noise-free digital (orange) trajectories. (*C*) The dwell time distributions of the ground-truth data. (*D*) The dwell time distributions generated from ebFRET extracted dwell times.

To ensure the ebFRET can robustly extract the dwell times, we compared the ebFRET extracted dwell time distributions with the ground-truth data. In short, using a 3-state kinetic model with a hidden state in the high state (Fig. S5*A*), we simulated single-molecule trajectories with given rate constants using a home-built software, SMIS (4). A random noise was then added onto the simulated trajectories to generate final trajectories that have similar signal-to-noise ratio as the experimental data (Fig. S5*B*). These simulated trajectories were then analyzed with ebFRET to extract the microscopic dwell times. Fig. S5B shows the overlay of the simulated (blue curve) and the ebFRET-extracted trajectories (orange curve) and demonstrate that the ebFRET faithfully captures the transitions. More importantly, the extracted dwell time distributions between states, including $\tau_{HM}$, $\tau_{MH}$, $\tau_{ML}$, and $\tau_{LM}$, (Fig. S5*C*), nicely reproduced the expected ground-truth data, validating the usage of ebFRET to get the microscopic dwell times for our experimental data (Fig. S5*D*).

### 7. Generation of probability density functions of dwell times

With the extracted microscopic dwell times from the cleaned-up SRF trajectories (Fig. S6*A*), we generated the probability density functions (*PDF*) of $\tau_{HM}$, $\tau_{MH}$, $\tau_{ML}$, and $\tau_{LM}$. Considering the first bin is often inaccurate due to limited time resolution, we removed the first bin in all dwell-time histograms (5). We further combined every two bins so that the final time resolution is 60 ms to ensure the histogram is statistically saturated (Fig. S6*B*). Fig. S6*C* summarizes $PDF_{HM}$, $PDF_{MH}$, $PDF_{ML}$, and $PDF_{LM}$ with AdoCbl concentration varying from 0 to 60 μM.

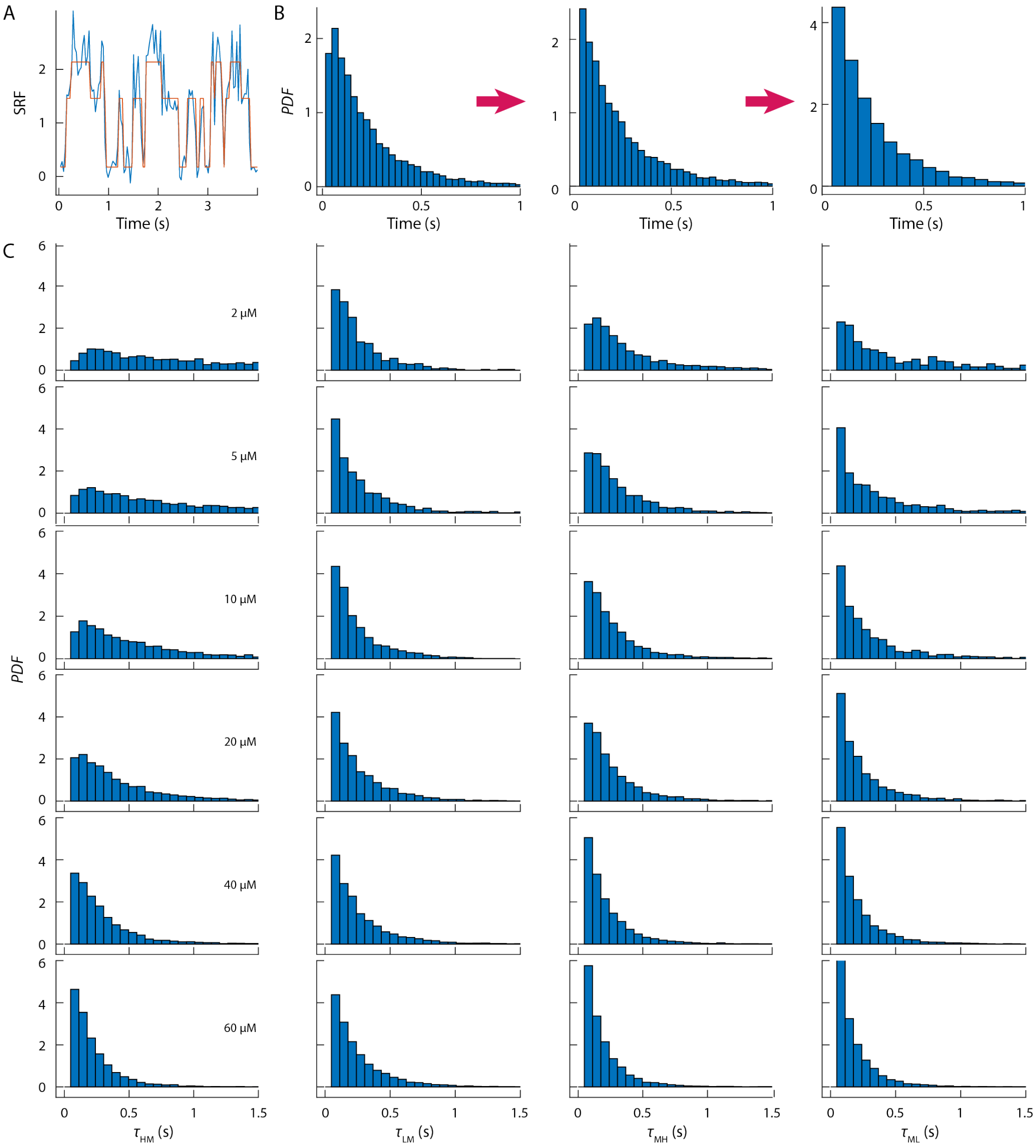


**Supplementary Fig. S6. Generation of probability density functions of dwell times.** (*A*) SRF trajectories (blue) were analyzed by ebFRET to extract the microscopic dwell times. (*B*) Dwell times from more than 200 SRF trajectories were used to generate the probability density function (*PDF*) (left). The first bin of the *PDF* was removed (middle) due to time-resolution issue. The remaining bins were further combined with 60 ms to ensure the data is statistically saturated (right). (*C*) *PDF*s of $\tau_{HM}$, $\tau_{LM}$, $\tau_{MH}$, and $\tau_{ML}$ with [AdoCbl] varies from 2 to 60 μM.

### 8. *MMAB*$_s$ is not the intermediate in the *F*$_H$ state

The exponential rise and decay feature of $PDF_{HM}$ at low [AdoCbl] indicates the presence of intermediate in the transition from $MMAB_U$ to $MMAB_S$. The characteristic signature requires the intermediate existing in the $F_H$ state. The simplest model is that the $MMAB_S$ in the $F_H$ state to be the intermediate. In this section, we tested this hypothesis by simulating the expected $PDF_{HM}$ and checking if this kinetical scheme reproduces the key observed kinetic features.

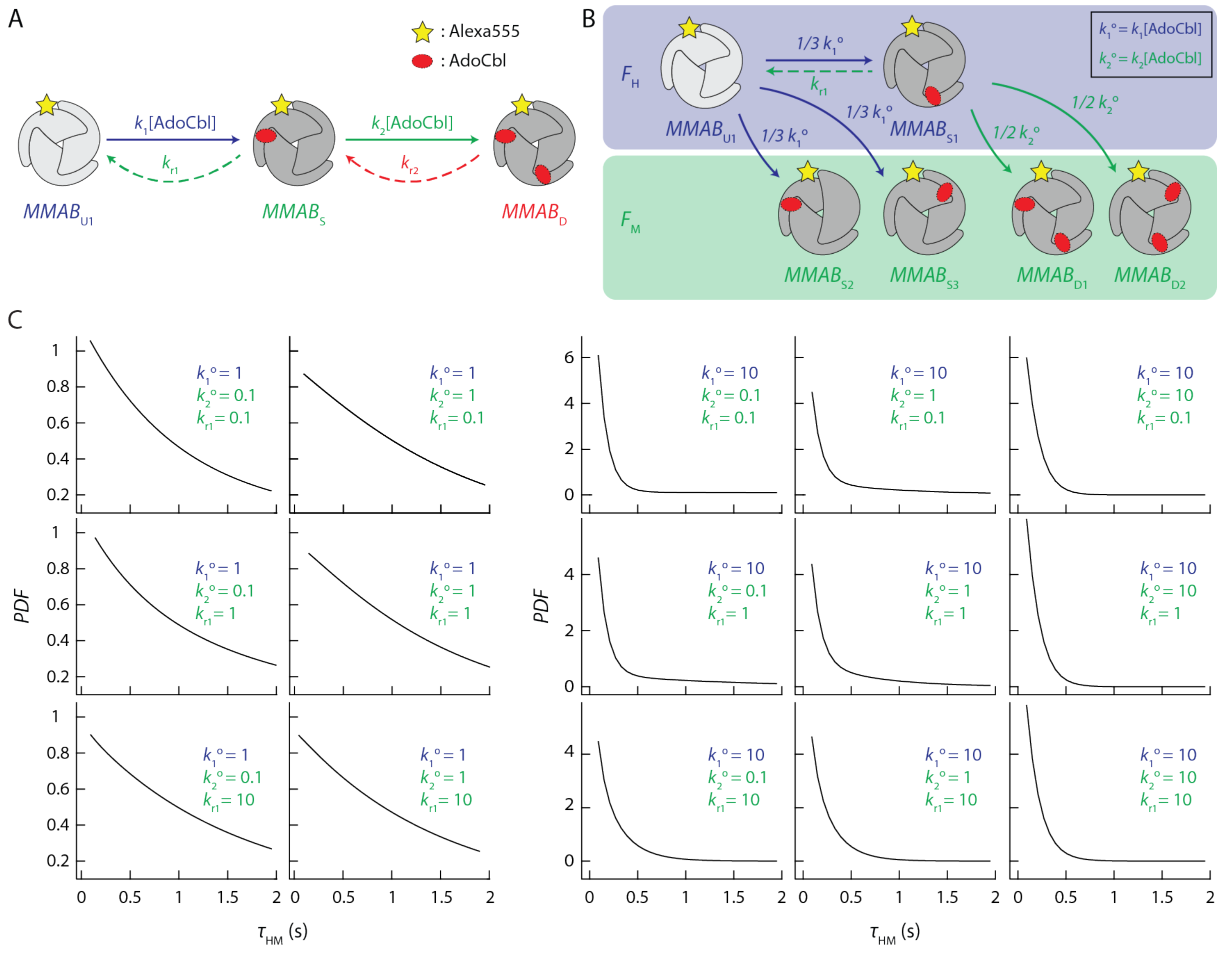

**Supplementary Fig. S7. *MMAB*s is not the intermediate in the *F*$_H$ state.** (*A*) The hypothesized kinetic model assuming the $MMAB_S$ is the intermediate in the $F_H$ state (top) and detailed submodel dictating $\tau_{HM}$ distributions (bottom). (*B*) Detailed kinetic model describing the transition steps between $F_H$ and $F_M$ states. (*C*) Simulations of $\tau_{HM}$ distributions based on the kinetic model cannot reproduce the exponential rise and decay feature of $PDF_{HM}$.

Using the kinetic model and the submodel dictating the different MMAB-AdoCbl binding configurations in Fig. S7*A*, we first solved the analytical solution of $\tau_{HM}$ distribution. The single-molecule rate equations related to the $\tau_{HM}$ distribution are:

$$\frac{dP_U(t)}{dt} = k_{-1}P_{S1}(t) - k_1^o P_U(t) \qquad S[1]$$

$$\frac{dP_{S1}(t)}{dt} = \frac{1}{3}k_1^o P_U(t) - (k_{-1} + k_2^o)P_{S1}(t) \qquad S[2]$$

$$\frac{dP_{S2}(t)}{dt} = \frac{dP_{S3}(t)}{dt} = \frac{1}{3}k_1^o P_U(t) \quad S[3]$$

$$\frac{dP_{D2}(t)}{dt} = \frac{dP_{D1}(t)}{dt} = \frac{1}{2}k_2^o P_{S1}(t) \quad S[4]$$

With initial condition: $P_U(0) = 1$, $P_{S1}(0) = P_{S2}(0) = P_{S3}(0) = P_{D1}(0) = P_{D2}(0) = 0$, and $P_U(t) + P_{S1}(t) + P_{S2}(t) + P_{S3}(t) + P_{D1}(t) + P_{D2}(t) = 1$, one can solve Eq S1- S4 and obtain

$$f_{HM}(\mathrm{t}) = \frac{k_1^o\left[(3k_2^o + 2k_{-1} + 2B_1 + 2A_1)e^{(B_1 + A_1)t} - (3k_2^o + 2k_{-1} + 2B_1 - 2A_1)e^{(B_1 - A_1)t}\right]}{6A_1} \quad S[5]$$

, where $A_1 = \frac{1}{2}\sqrt{k_1^{o2} + (k_2^o + k_{-1})^2 - \frac{2}{3}k_1^o(3k_2^o + k_{-1})}$ and $B_1 = -\frac{1}{2}(k_1^o + k_2^o + k_{-1})$.

We then simulated the expected $PDF_{HM}$ with the analytical solution of $\tau_{HM}$ to examine whether the hypothesized kinetic model can recapture the exponential rise and decay feature of $PDF_{HM}$ at low [AdoCbl]. It is worth noting that Fig 3*A* shows the reaction rate of exiting $F_H$ to $F_M$ state is dependent on [AdoCbl]. The steeper slope in the low [AdoCbl] region indicates the formation rate constant of $MMAB_S$ is larger than that of $MMAB_D$ (e.g., $k_1 > k_2$). Using this criterion, we simulated the $PDF_{HM}$ with $k_1^o$(i.e., $k_1$[AdoCbl]), $k_{-1}$, $k_2^o$(i.e., $k_2$[AdoCbl]) with 0.1, 1, 10 $s^{-1}$ (Fig. S7*B*). All simulations show simple decay curves and fail to reproduce the exponential rise and decay feature, excluding the $MMAB_S$ in the $F_H$ state from the possible intermediate. We thus propose that the intermediate originates from the unbound MMAB under a different conformation ($MMAB_{U2}$).

### 9. *MMAB*$_S$ must return to *MMAB*$_{U1}$ when AdoCbl dissociation occurs

With the presence of *MMAB*$_{U2}$, dissociations of AdoCbl from the *MMAB*$_S$ can either go to *MMAB*$_{U1}$ or *MMAB*$_{U2}$. To identify the correct pathway, we hypothesized that *MMAB*$_{U1}$ is the destination and checked if the kinetic model (Fig. S8*A*) could reproduce the experimental observations.

Using the $k_{r1}$ and $k_2$ from $PDF_{LMH}$ and $PDF_{LML}$ (details in section 10), we use SMIS to simulate the $PDF_{HM}$ based on the Fig. S8A. We varied $k^i_1$, $k^i_{-1}$, and $k^o_1$ 0.1 to 10 to test if any combination could reproduce the unique exponential rise and decay feature. Fig. S8*B-D* show that all combinations give single-exponentials decay curves. This discrepancy between simulation and experimental results confirmed that *MMAB*$_S$ doesn't return to *MMAB*$_{U2}$ when AdoCbl dissociates. We thus finalize our kinetic model that *MMAB*$_S$ must return to *MMAB*$_{U1}$ when AdoCbl dissociates (Fig. 3*G*).

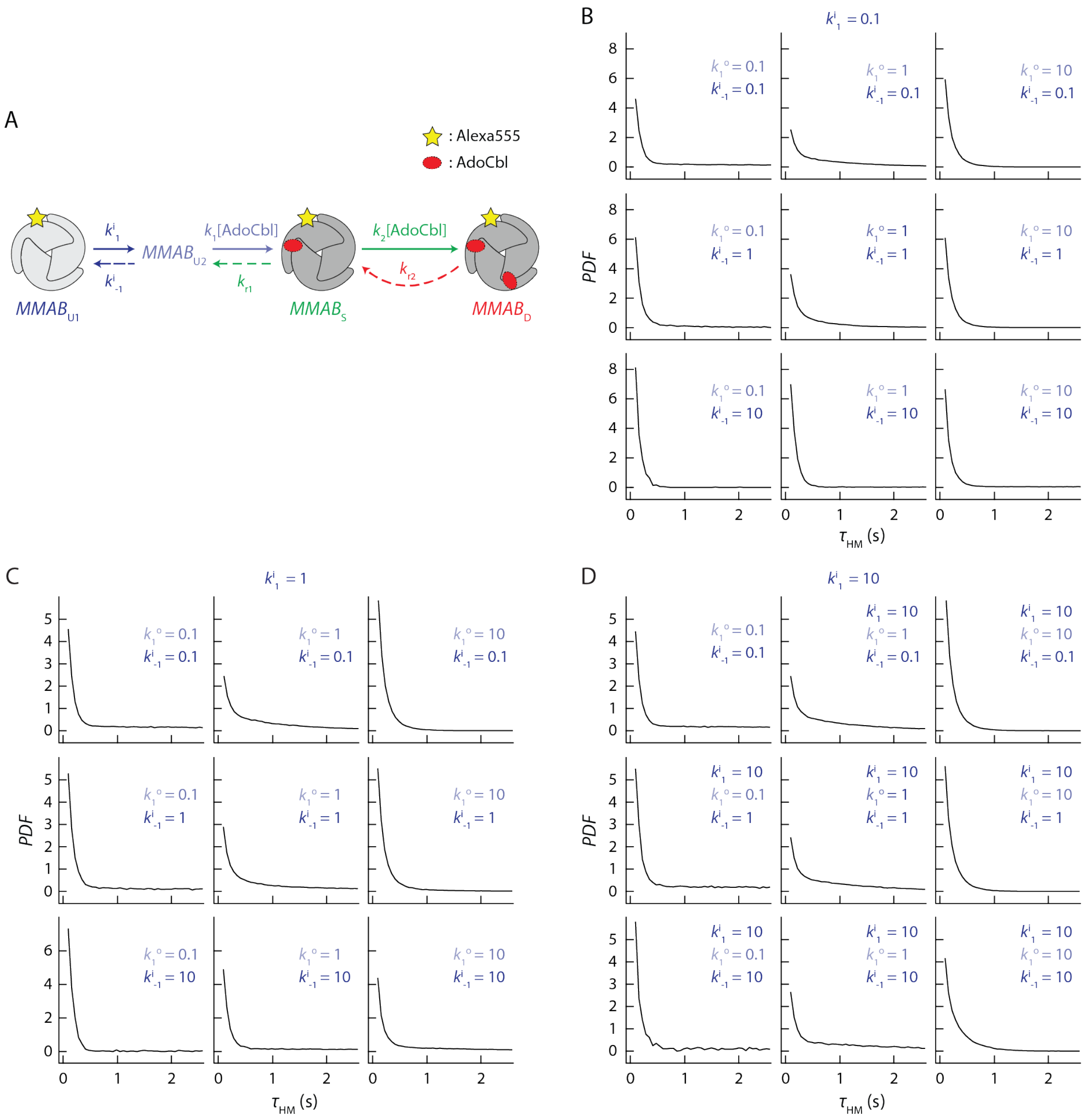


**Supplementary Fig. S8. *MMAB*$_S$ must return to *MMAB*$_{U1}$ when AdoCbl dissociation occurs.** (*A*) The hypothesized kinetic model assuming the dissociation of AdoCbl from *MMAB*$_S$ resulted the formation of *MMAB*$_{U2}$. (*B-D*) Simulations of $\tau_{HM}$ distributions based on the kinetic model in *A* cannot reproduce the exponential rise and decay feature of $PDF_{HM}$.

### 10. Derivations of $f_{LMH}(\tau)$ and $f_{LML}(\tau)$

We derived the analytical expression of the probability density function of dwell time, $f_{LMH}(\tau)$ and $f_{LML}(\tau)$, using the kinetic model presented in Fig. S9. MMAB with different configurations were labeled individually. For example, the $MMAB_S$ in the $F_H$ was labeled as $MMAB_{S1}$, and the remaining $MMAB_S$ in the $F_M$ were labeled as $MMAB_{S2}$ and $MMAB_{S3}$. The two $MMAB_D$ in the $F_M$ were labeled as $MMAB_{D1}$ and $MMAB_{D2}$, and the remaining $MMAB_D$ in the $F_L$ was labeled as $MMAB_{D3}$. Note that the following species: (1) $MMAB_{S2}$ and $MMAB_{S3}$, (2) $MMAB_{D1}$ and $MMAB_{D2}$ are two equivalent and undistinguishable pairs in our model. We, therefore, combined each group to simplify the derivations.

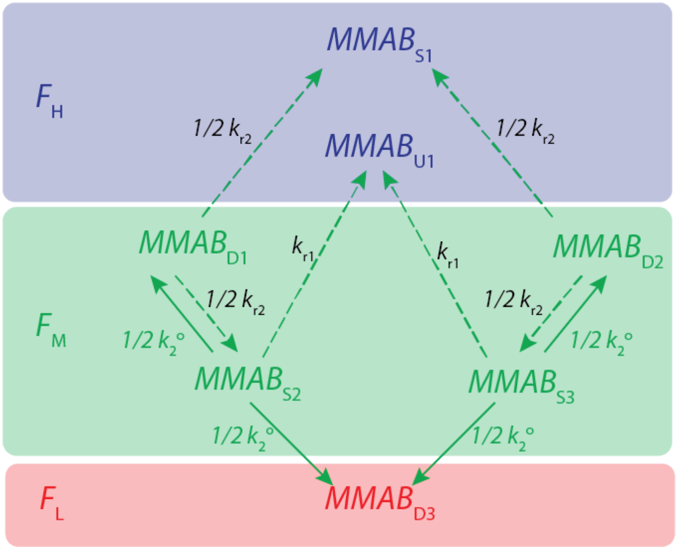


**Supplementary Fig. S9. The kinetic scheme highlighted all processes associated with $\tau_{LML}$ and $\tau_{LMH}$.**

To solve $f_{LMH}(\tau)$ and $f_{LML}(\tau)$, we started with the analytical expression of the probabilities of finding the species $MMAB_{S3}$, $P_{S3}(t)$. We first focused on the steps that involve the transition $P_{S2}(t)$ and $P_{S3}(t)$. The single-molecule rate equations for these steps are:

$$\frac{dP_{S3}(t)}{dt} = \frac{dP_{S2}(t)}{dt} = \frac{1}{2}k_{r2}P_{D1}(t) - (k_2^0 + k_{r1})P_{S3}(t) \qquad S[6]$$

$$\frac{dP_{D1}(t)}{dt} = \frac{dP_{D2}(t)}{dt} = \frac{1}{2}k_2^0P_{S3}(t) - k_{r2}P_{D1}(t) \qquad S[7]$$

, where the $P_i(t)$ are the probabilities of finding the species ($i$ = $S_2$, $S_3$, $D_2$, $D_3$) in the corresponding states at time $t$. Note that $k_2^0$ (i.e., $k_2$[S]) is treated as pseudo-first-order rate constants and $P_i'(t)$ is equal to $dP_i(t)/dt$. We first applied Laplace transform to Eq. S6 and Eq. S7:

$$\mathcal{L}\{P_{S3}'(t)\} = \frac{1}{2}k_{r2}\mathcal{L}\{P_{D1}(t)\} - (k_2^0+k_{r1})\mathcal{L}\{P_{S3}(t)\} \qquad S[8]$$

$$\mathcal{L}\{P_{D1}'(t)\} = \frac{1}{2}k_2^0\mathcal{L}\{P_{S3}(t)\} - k_{r2}\mathcal{L}\{P_{D1}(t)\} \qquad S[9]$$

Each $\mathcal{L}\{P_i'(t)\}$ in Eq. S8 and Eq. S9 were equal to $s\mathcal{L}\{P_i(t)\}$ - $P_i(0)$. Since we defined all transitions starting from $MMAB_{S2}$ or $MMAB_{S3}$, we obtained the following boundary conditions: $P_{S3}(0)$ = $P_{S3}(0)$ = 0.5, $P_{D1}(0)$ =

$P_{D2}(0) = 0$. By substituting $\mathcal{L}\{P_i'(t)\}$ with $s\mathcal{L}\{P_i(t)\}$ - $P_i(0)$ in Eq. S8 and Eq. S9 and replaced $P_i(0)$ with values specified in the boundary conditions, we expressed $\mathcal{L}\{P_i'(t)\}$ as:

$$(s + k_2^o + k_{r1})\mathcal{L}\{P_{S3}(t)\} - 0.5 = \frac{1}{2}k_{r2}\mathcal{L}\{P_{D1}(t)\} \quad S[10]$$

$$\mathcal{L}\{P_{D1}(t)\} = \frac{1}{2}k_2^0\mathcal{L}\{P_{S3}(t)\}/(s + k_{r2}) \quad S[11]$$

Substituting $\mathcal{L}\{P_{D1}(t)\}$ in Eq. S10 using Eq. S11, we can express $\mathcal{L}\{P_{S3}(t)\}$ with rate constants as:

$$\mathcal{L}\{P_{S3}(t)\} = \frac{b + k_{r2}}{2(\frac{1}{4}k_2^o k_{r2} - (b + k_{r2})(b + k_2^o + k_{r2}))} \quad S[12]$$

To derive the $f_{LMH}(\tau)$ in the two-state kinetic model, we focused on the steps involve the transition from $MMAB_{S2}$ or $MMAB_{S3}$ → $MMAB_{U1}$ and $MMAB_{D1}$ or $MMAB_{D2}$ → $MMAB_{S1}$ (Fig. S9). The single-molecule rate equations for these steps are:

$$\frac{dP_{U1}(t)}{dt} = k_{r1}P_{S3}(t) + k_{r1}P_{S2}(t) = 2k_{r1}P_{S3}(t) \quad S[13]$$

$$\frac{dP_{S1}(t)}{dt} = \frac{1}{2}k_{r2}P_{D1}(t) + \frac{1}{2}k_{r2}P_{D2}(t) = k_{r2}P_{D1}(t) \quad S[14]$$

Here, the initial conditions are $P_{U1}(0) = 0$ and $P_{S1}(0) = 0$ at $t = 0$. Take the Laplace transform of Eq. S13 and Eq. S14, we obtained:

$$\mathcal{L}\{P_{U1}'(t)\} = 2k_{r1}\mathcal{L}\{P_{S3}(t)\} \quad S[15]$$

$$\mathcal{L}\{P_{S1}'(t)\} = k_{r2}\mathcal{L}\{P_{D1}(t)\} \quad S[16]$$

$\mathcal{L}\{P_{D1}(t)\}$ in Eq. S16 was further replaced by Eq. S11, which resulted in a new $\mathcal{L}\{P_{S1}'(t)\}$ expressed as a function of $\mathcal{L}\{P_{S3}(t)\}$:

$$\mathcal{L}\{P_{S1}'(t)\} = \frac{k_{r2}k_2^0\mathcal{L}\{P_{S3}(t)\}}{2(s + k_{r2})} \quad S[17]$$

$\tau_{LMH}$ is the time needed to complete steps involving $k_2$, $k_{r1}$ and $k_{r2}$. The probability of finding a particular $\tau$ is equal to the probability for the switch from $MMAB_{S2}$ or $MMAB_{S3}$ → $MMAB_U$ and $MMAB_{D1}$ or $MMAB_{D2}$ → $MMAB_{S1}$ state between $t = \tau$ and $\tau + \Delta\tau$. Mathematically, this probability is equal to $\Delta P_{S1}(\tau) + \Delta P_{U1}(\tau)$. In the limit of infinitesimal $\Delta\tau$, the probability density function of dwell time $\tau_{LMH}$, $f_{LMH}(t)$, was obtained from

$$f_{LMH}(t) = \frac{dP_{S1}(t)}{dt} + \frac{dP_{U1}(t)}{dt} = P_{S1}'(t) + P_{U1}'(t) \quad S[18]$$

Take the Laplace transform of Eq. S18, we obtained:

$$\mathcal{L}\{f_{LMH}(t)\} = \mathcal{L}\{P_{S1}'(t) + P_{U1}'(t)\} = \mathcal{L}\{P_{S1}'(t)\} + \mathcal{L}\{P_{U1}'(t)\} \quad S[19]$$

Substituting the $\mathcal{L}\{P'_{S1}(t)\}$ and $\mathcal{L}\{P'_{U1}(t)\}$ in Eq. S19 with Eq. S15 and Eq. S17; replacing $\mathcal{L}\{P_{S3}(t)\}$ using Eq. S12, we got:

$$\mathcal{L}\{f_{LMH}(t)\} = \frac{k_{r1}s + (\frac{1}{4}k_2^0 k_{r2} + k_{r1}k_{r2})}{s^2 + (k_2^0 + k_{r1} + k_{r2})s + (\frac{3}{4}k_2^0 k_{r2} + k_{r1}k_{r2})} \qquad S[20]$$

Inverse Laplace transform of Eq. S20 gave the probability density function of dwell time, $f_{LMH}(\tau)$:

$$f_{LMH}(\tau) = \frac{4k_{r1}(b + a + k_{r2}) + k_2^0 k_{r1}}{8a} e^{(b+a)\tau} - \frac{4k_{r1}(b - a + k_{r2}) + k_2^0 k_{r1}}{8a} e^{(b-a)\tau} \qquad S[21]$$

, where $a = \frac{1}{2}\sqrt{k_2^{0\,2} + k_{r1}^{\ 2} + k_{r2}^{\ 2} + 2k_2^0 k_{r1} - k_2^0 k_{r2} - 2k_{r1}k_{r2}} = \sqrt{b^2 - \frac{3}{4}k_2^0 k_{r2} - k_{r1}k_{r2}}$ and $b = -\frac{1}{2}(k_2^0 + k_{r1} + k_{r2})$

To derive the $f_{LML}(\tau)$ in the two-state kinetic model, we focused on the steps involve the transition to $MMAB_D$ (Fig. S9). The single-molecule rate equation for these steps is:

$$\frac{dP_{D3}(t)}{dt} = \frac{1}{2}k_2^0 P_{S2}(t) + \frac{1}{2}k_2^0 P_{S3}(t) = k_2^0 P_{S3}(t) \qquad S[22]$$

Here, the initial condition is $P_{D3}(0) = 0$ at $t = 0$. Take the Laplace transform of Eq. S22, we obtained:

$$\mathcal{L}\{P'_{D3}(t)\} = k_2^0 \mathcal{L}\{P_{S3}(t)\} \qquad S[23]$$

$\tau_{LML}$ is the time needed to complete steps involving $k_2$, $k_{r1}$ and $k_{r2}$. The probability of finding a particular $\tau$ is equal to the probability for the switch from $MMAB_{S3}$ or $MMAB_{S3} \rightarrow MMAB_{D3}$ between $t = \tau$ and $\tau + \Delta\tau$. Mathematically, this probability is equal to $\Delta P_{D3}(\tau)$. In the limit of infinitesimal $\Delta\tau$, the probability density function of dwell time $\tau_{LML}$, $f_{LML}(t)$, was obtained from

$$f_{LML}(\tau) = \frac{dP_{D3}(\tau)}{d\tau} = P'_{D3}(\tau) \qquad S[24]$$

Take the Laplace transform of Eq. S24, we obtained:

$$\mathcal{L}\{f_{LML}(\tau)\} = \mathcal{L}\{P'_{D3}(\tau)\} \qquad S[25]$$

Substituting the $\mathcal{L}\{P'_{D3}(\tau)\}$ in Eq. S25 with Eq. S23; replacing $\mathcal{L}\{P_{S3}(t)\}$ using Eq. S12, we got:

$$\mathcal{L}\{f_{LML}(\tau)\} = \frac{\frac{1}{2}k_2^0(s + k_{r2})}{s^2 + (k_2^0 + k_{r1} + k_{r2})s + (\frac{3}{4}k_2^0 k_{r2} + k_{r1}k_{r2})} \qquad S[26]$$

Inverse Laplace transform of Eq. S26 gave the probability density function of dwell time, $f_{LML}(\tau)$:

$$f_{LML}(\tau) = \frac{k_2^0}{4a}[(b + a + k_{r2})\, e^{(b+a)\tau} - (b - a + k_{r2})e^{(b-a)\tau}] \qquad S[27]$$

, where $a$ and $b$ are defined the same as in Eq. S21. Finally, the analytical probability density functions, $f_{LML}(\tau)$ and $f_{LMH}(\tau)$, were used to global fitting the distributions of $\tau_{LML}$ and $\tau_{LMH}$ by MATLAB and extracted out the rate constants $k_2$, $k_{r1}$ and $k_{r2}$.

### 11. Derivation of $K_{D1}$ and $K_{D2}$

Using the proposed kinetic model, we derived the analytical solution of the equilibrium dissociation constants for $MMAB_S$ and $MMAB_D$ from the population relationship. Assuming the system is under dynamic equilibrium condition, the populations of $MMAB_{U1}$ ($P_{U1}$), $MMAB_{U2}$ ($P_{U2}$), $MMAB_S$ ($P_S$) and $MMAB_D$ ($P_D$) can be associated with rate constants and [AdoCbl] ($L$) through equations below:

$$k_{r1}P_S + k^i_{-1}P_{U2} = k^i_1 P_{U1} \quad \text{S[28]}$$

$$k_1 L P_{U2} + k_{r2}P_D = (k_{r1} + k_2 L)P_S \quad \text{S[29]}$$

$$k_2 L P_S = k_{r2}P_D \quad \text{S[30]}$$

$$k^i_1 P_{U1} = (k_1 L + k^i_{-1})P_{U2} \quad \text{S[31]}$$

$$P_{U1} + P_S + P_D + P_{U2} = 1 \quad \text{S[32]}$$

By solving Eq. S28-32, we expressed the populations of these species as a function of rate constants as below:

$$P_{U1} = \frac{(k_1 L + k^i_{-1})\, k_{r1}\, k_{r2}}{C} \quad \text{S[33]}$$

$$P_S = \frac{k^i_1 k_1 k_{r2} L}{C} \quad \text{S[34]}$$

$$P_D = \frac{k^i_1 k_1 k_2 L^2}{\mathrm{C}} \quad \text{S[35]}$$

$$P_{U2} = \frac{k^i_1 k_{r1} k_{r2}}{C} \quad \text{S[36]}$$

, where $C = k^i_1 k_{r1} k_{r2} + k_{r1} k_{r2}(k^i_{-1} + k_1 L) + k^i_1 k_1 \mathrm{L}(k_{r2} + k_2 L)$.

We then defined the fractional occupancy $\Omega(L)$ as the population of the bound species over the population of all species that are involved in the equilibrium, so we can get the expression of the first binding fractional occupancy $\Omega_1(L)$ and the second binding fractional occupancy $\Omega_2(L)$ as following:

$$\Omega_1(L) = \frac{P_S}{P_{U1} + P_{U2} + P_S} \quad \text{S[37]}$$

$$\Omega_2(L) = \frac{P_D}{P_S + P_D} \quad \text{S[38]}$$

Substituting $P_{U1}$, $P_S$, $P_D$, and $P_{U2}$ in Eq. S37- 38 with Eq. S33- 36, we got:

$$\Omega_1(L) = \frac{k^i_1 k_1 L}{k_{r1}(k^i_{-1} + k_1 L) + k^i_1(k_{r1} + k_1 L)} \quad \text{S[39]}$$

$$\Omega_2(L) = \frac{k_2 L}{k_2 L + k_{r2}} \qquad \text{S[40]}$$

We then defined the $K_{D1}$ and $K_{D2}$ (5) by the concentration of AdoCbl at which the $\Omega_1(L) = \frac{1}{2} \times \Omega_{1,max}(L)$ and $\Omega_2(L) = \frac{1}{2} \times \Omega_{2,max}(L)$, respectively:

$$K_{D1} = \frac{k_{r1}(k_1^i + k_{-1}^i)}{k_1(k_1^i + k_{r1})} \qquad \text{S[41]}$$

$$K_{D2} = \frac{k_{r2}}{k_2} \qquad \text{S[42]}$$

Eq. S41 and S42 can be expressed in terms of the overall forward (i.e., $k_{f1}$ and $k_{f2}$) and reverse rate constants (i.e., $k_{r1}$ and $k_{r2}$) where $k_{f1} = k_1(k_1^i+k_{r1})/(k_1^i+k_{-1}^i)$.

$$K_{D1} = \frac{k_{r1}(k_1^i + k_{-1}^i)}{k_{f1}} = \frac{k_{r1}}{k_1(k_1^i + k_{r1})/(k_1^i + k_{-1}^i)} \qquad \text{S[43]}$$

$$K_{D2} = \frac{k_{r2}}{k_{f2}} = \frac{k_{r2}}{k_2} \qquad \text{S[44]}$$

Using the rate constants listed in Table 1, we estimate the $K_{D1,\ Ado}$ = 5.6 ± 2.9 µM and $K_{D2,\ Ado}$ = 41.7 ± 13.9 µM. $K_{D2,\ Ado}$ is ~7 times larger than $K_{D1,\ Ado}$, supporting the negative cooperative binding of AdoCbl to MMAB. The overall reverse rate constant of $MMAB_S$ ($k_{r1}$) and $MMAB_D$ ($k_{r1}$) are 3.7 $s^{-1}$ and 5.0 $s^{-1}$, respectively. $k_{r2}$ is 1.35 times of $k_{r1}$, which suggests the dissociation processes are not the dominant contributors to the negative cooperativity. While the overall forward rate constant of $MMAB_{U1}$ ($k_{f1}$) and $MMAB_S$ ($k_{f2}$) are 0.66 $s^{-1}$ and 0.12 $s^{-1}$, respectively. $k_{f1}$ is 5.5 times higher than $k_{f2}$, which suggests the association process is the major factor that causes negative cooperativity. We also estimate $K_{D1,\ OHCbl}$ = 15.0 ± 5.0 µM and $K_{D2,\ OHCbl}$ = 189 ± 111 µM. $K_{D2,\ OHCbl}$ is ~ 13 times larger than $K_{D1,\ OHCbl}$. However, the $K_{D2,\ OHCbl}$ is beyond our experimental [OHCbl] and has large error bar, making the effect of second OHCbl binding unclear.

### 12. SRF state assignments and subpopulation analysis for MMAB-OHCbl interactions

Similar experiments and analyses were performed to study MMAB-OHCbl interaction kinetics. But due to the weaker binding affinity between MMAB and OHCbl ($K_D$ ~ 200 µM) (6), we varied the [OHCbl] concentration from 0 to 80 µM. Similar to the AdoCbl experiments, three fluorescence states, $F_H$, $F_M$, and $F_L$, were observed. As [OHCbl] increases, the $F_M$ (~ 0.6) and $F_L$ (~ 0.1) peaks gradually appear while the $F_H$ peak decreases (Fig. 5*A*). We globally fit the normalized SRF intensity histograms across all [OHCbl] with a three-Gaussian model. The center position and width of each Gaussian distribution were shared across all [OHCbl] while keeping the relative populations among three Gaussians to float. Fig. 5*B* summarizes the fitted subpopulations, which were used to associate different MMAB-OHCbl binding configurations (Fig. 5C) with the $F_H$, $F_M$, and $F_L$ states.

In the MMAB-AdoCbl experiments, we analyze the subpopulations of each SRF and summarized the SRF assignments in Fig. 2C. The $MMAB_U$ and the $MMAB_D$ with two AdoCbl bound at the near two sites ($MMAB_{D3}$) undoubtedly contribute to the $F_H$ and $F_L$ states, respectively. The singly bound MMAB, $MMAB_S$, previously were grouped into two subgroups in the MMAB-AdoCbl study. The MMAB with AdoCbl bound at the near two sites ($MMAB_{S2}$ and $MMAB_{S3}$) shows clear quenching and assigned to the $F_M$ state, while at the farthest site assigned to the $F_H$ state. However, the absorption spectrum of cobalamin and the Alexa555 emission spectrum (Fig. 1*A*) indicates that the OHCbl is a stronger quencher than the AdoCbl due to its larger spectral overlap with the Alexa555 dye. The stronger FRET efficiency of OHCbl may result in new fluorescence state assignment towards the lower fluorescence states. In other words, $MMAB_S$ and $MMAB_D$ with a OHCbl bound at the farthest site ($MMAB_{D1}$ and $MMAB_{D2}$) may need to be assigned to the $F_M$ and the $F_L$ states, respectively.

To test this possibility, we checked the subpopulations of SRF states and compared various assignments (Table S3). At the highest substrate concentration (i.e., [OHCbl] = 80 µM), the subpopulation of the $F_H$, $F_M$, and $F_L$ is 37%, 57%, and 6%, respectively (Fig. 5C). We assume that $MMAB_s$ with OHCbl bound at the farthest site ($MMAB_{S1}$) does not increase the quenching and remains in the $F_H$ state (Table S3, assignment 1). In this case, the state assignment will be the same as the MMAB-AdoCbl. The $MMAB_D$ overall population will be 18% (6% from $F_L$ and 12% from $F_M$); the $MMAB_S$ overall population will be 67.5% (45% from $F_M$ and 22.5% from $F_H$); and The $MMAB_U$ overall population will be 14.5%. Since each MMAB can maximumly bind 2 OHCbl, we can estimate the fraction of site occupancy using the relative populations of $MMAB_U$, $MMAB_S$, and $MMAB_D$. Assuming we have 100 MMAB molecules (i.e., 200 available binding sites), assignment 1 gives a total 171 sites occupied by the OHCbl, leading to a final percent

**Table S3. SRF state assignments for MMAB-OHCbl complexes**

| SRF state | population (80 µM) | assignment 1 | assignment 2 | assignment 3 |
|---|---|---|---|---|
| $F_H$ | 37% | $MMAB_U$ (14.5%)<br>$MMAB_{S1}$(22.5%) | $MMAB_U$ | $MMAB_U$<br>37% |
| $F_M$ | 57% | $MMAB_{S2}$(22.5%), $MMAB_{S3}$(22.5%)<br>$MMAB_{D1}$(6%), $MMAB_{D2}$(6%) | $MMAB_{S1}$ | $MMAB_{S1}$, $MMAB_{S2}$, $MMAB_{S3}$,<br>57% |
| $F_L$ | 6% | $MMAB_{D3}$(6%) | $MMAB_{S2}$, $MMAB_{S3}$,<br>$MMAB_{D1}$, MMAB$_{D2}$,<br>$MMAB_{D3}$ | $MMAB_{D1}$, MMAB$_{D2}$, $MMAB_{D3}$<br>6% |
| **site occu.**[a] | | 51.75% | | 35% |
| **app. $K_D$**[b] | | < 80 µM | | > 80 µM |

[a] site occupancy. [b] apparent $K_D$.

occupancy of 51.75% (i.e., 103.5/200) at [OHCbl] of 80 µM. This means the apparent $K_D$ will be below 80 µM, which is in contradictory with the reported $K_D$ of 200 µM. In other words, $MMAB_{S1}$ can't be assigned to the $F_H$ state. Since $MMAB_{S1}$ is the configuration with the weakest quenching efficiency, we thus concluded that the MMAB with OHCbl bound at the farthest site must be reassigned to the $F_M$ state.

We then focus on the SRF assignment of the other two singly bound MMAB, $MMAB_{S2}$ and $MMAB_{S3}$. Assuming the binding of OHCbl shift the $MMAB_{S2}$ and $MMAB_{S3}$ to the $F_L$ state, the doubly bound MMAB, $MMAB_{D1}$ and $MMAB_{D2}$ must also shift to the $F_L$ state, which results in assignment 2. However, this specific assignment left $MMAB_{S1}$ as the only species in the $F_M$ state with a population of 57%. Since the overall population of $F_L$ state is only 6%, it is impossible to calculate the population of $MMAB_{S2}$ and $MMAB_{S3}$. We thus concluded that $MMAB_{S2}$ and $MMAB_{S3}$ do not contribute to the $F_L$ state. This information collectively suggests that $MMAB_{S1}$, $MMAB_{S2}$, and $MMAB_{S3}$ belonged to the $F_M$ state.

The fact that $MMAB_{S1}$, $MMAB_{S2}$, and $MMAB_{S3}$ are assigned to the same SRF state indicates that the quenching efficiency at three binding sites are comparable. This also implies that all the doubly bound MMAB should be reassigned to the $F_L$ state (assignment 3). This assignment predicts the site occupancy of 35% and apparent $K_d$ larger than 80 µM, agreeing well with the literature reported values. We thus finalized the SRF assignment with $MMAB_U$, $MMAB_S$, and $MMAB_D$ existing in the $F_H$, $F_M$, and $F_L$ states, respectively.

### 13. Minimal kinetic model for MMAB-OHCbl interactions

$\tau_{HM}$ and $\langle\tau_{HM}\rangle^{-1,}$ respectively, represent the dwell time and the averaged single-molecule transition rate of transitions from the $F_H$ state to the $F_M$ state. Both contain information on the formation of $MMAB_{S,}$ depending on the [OHCbl]. Fig. S10*A* shows that $\langle\tau_{HM}\rangle^{-1}$ linearly increases with increasing [OHCbl], which is expected for the OHCbl binding. The distribution of $\tau_{HM}$, instead of a simple single-exponential decay, shows an exponential rise and decay up to [OHCbl] = 40 µM (Fig. S10*B*) similar to the AdoCbl case. The delayed maximum at $\tau_{HM} > 0$ indicates the presence of the intermediate. Since the unbound MMAB, $MMAB_{U1}$, is the only configuration that contribute to the $F_H$ state, we thus propose the intermediate a new MMAB conformation, $MMAB_{U3}$, to bind the OHCbl.

$\tau_{LM}$ and $\langle\tau_{LM}\rangle^{-1}$ are parameters relative to OHCbl unbinding behaviors from the $MMAB_D$. The distribution of $\tau_{LM}$ can be well-fitted with a single exponential decay to report the dissociation rate constants. Both $\langle\tau_{LM}\rangle^{-1}$ and fitted rate constants remain the same across all [OHCbl] (Fig. S10*C*). This [OHCbl] independent rate constant indicates the unbinding of OHCbl from the $MMAB_D$ through the typical unimolecular unbinding mechanism.

With the presence of $MMAB_{U3}$, dissociations of OHCbl from the $MMAB_S$ can either go to $MMAB_{U1}$ or $MMAB_{U3}$. To identify the correct pathway, we hypothesized that $MMAB_{U3}$ is the destination and checked if the kinetic model (Fig. S10*D*) could reproduce the experimental observations. Using the $k_{r1}$ and $k_2$ from $PDF_{MH}$ and $PDF_{ML}$ and $k_{r2}$ from $PDF_{LM}$ (details in section 14), we used SMIS to simulat the $PDF_{HM}$ based on the Fig. S11A. We varied $k_1^i$, $k_{-1}^i$, and $k_1^o$ 0.1 to 10 to test if any combination could reproduce the unique exponential rise and decay feature. Fig. S10*E-G* show that all combinations give single-exponentials decay curves. This discrepancy between simulation and experimental results confirmed that $MMAB_S$ doesn't return to $MMAB_{U3}$ when OHCbl dissociates. We thus finalize our kinetic model that $MMAB_S$ must return to $MMAB_{U1}$ when OHCbl dissociates.

With the above analysis, we proposed a minimal kinetic model that can describe our observations for MMAB-OHCbl interactions. The binding of OHCbl to MMAB shows clear [OHCbl] dependence. The binding of the first OHCbl requires the formation of intermediate (i.e., the delayed maximum at $\tau_{HM} > 0$ across all low [OHCbl]); Dissociation of OHCbl from $MMAB_S$ and $MMAB_D$ follows unimolecular dissociation mechanism (i.e., [OHCbl] independent dissociation rate), with the destination specie to be $MMAB_{U1}$ (similar to the AdoCbl case in section S9). Combining this information, we formulated the minimal kinetic model of MMAB-OHCbl interactions (Fig. 5*E*).

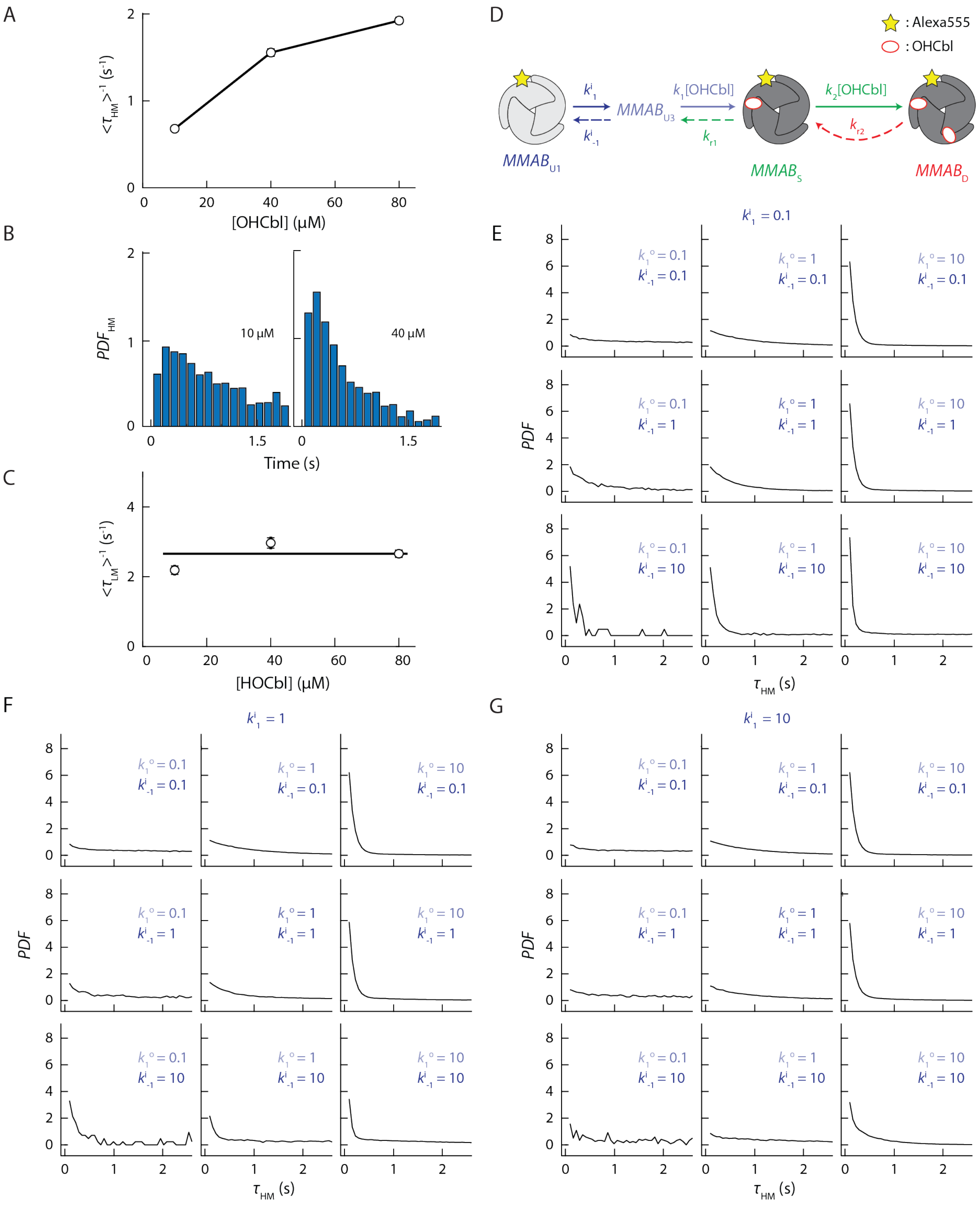


**Supplementary Fig. S10. Dwell time analysis and minimal kinetic model of MMAB-OHCbl interactions.** (*A*) [OHCbl] dependent $<\tau_{HM}>^{-1}$ shows the averaged binding rate increases with higher OHCbl concentration. (*B*) Probability density functions of $\tau_{HM}$ under low and high OHCbl concentrations. (*C*) [OHCbl] dependent $<\tau_{LM}>^{-1}$ shows the unbinding rate of HOCbl from $MMAB_D$ is [OHCbl] independent. (*D*) The hypothesized kinetic model assuming the dissociation of OHCbl from $MMAB_S$ resulted the formation of $MMAB_{U3}$. (*E-G*) Simulations of $\tau_{HM}$ distributions based on the kinetic model in *D* cannot reproduce the exponential rise and decay feature of $PDF_{HM}$.

### 14. Extraction of kinetic rate constants for MMAB-OHCbl interactions

To extract the kinetic parameters for MMAB-OHCbl interactions, we derived analytical dwell time solutions to dissect kinetic rate constants for MMAB-OHCbl interactions based on the minimal kinetic model (Fig. 5*E*).

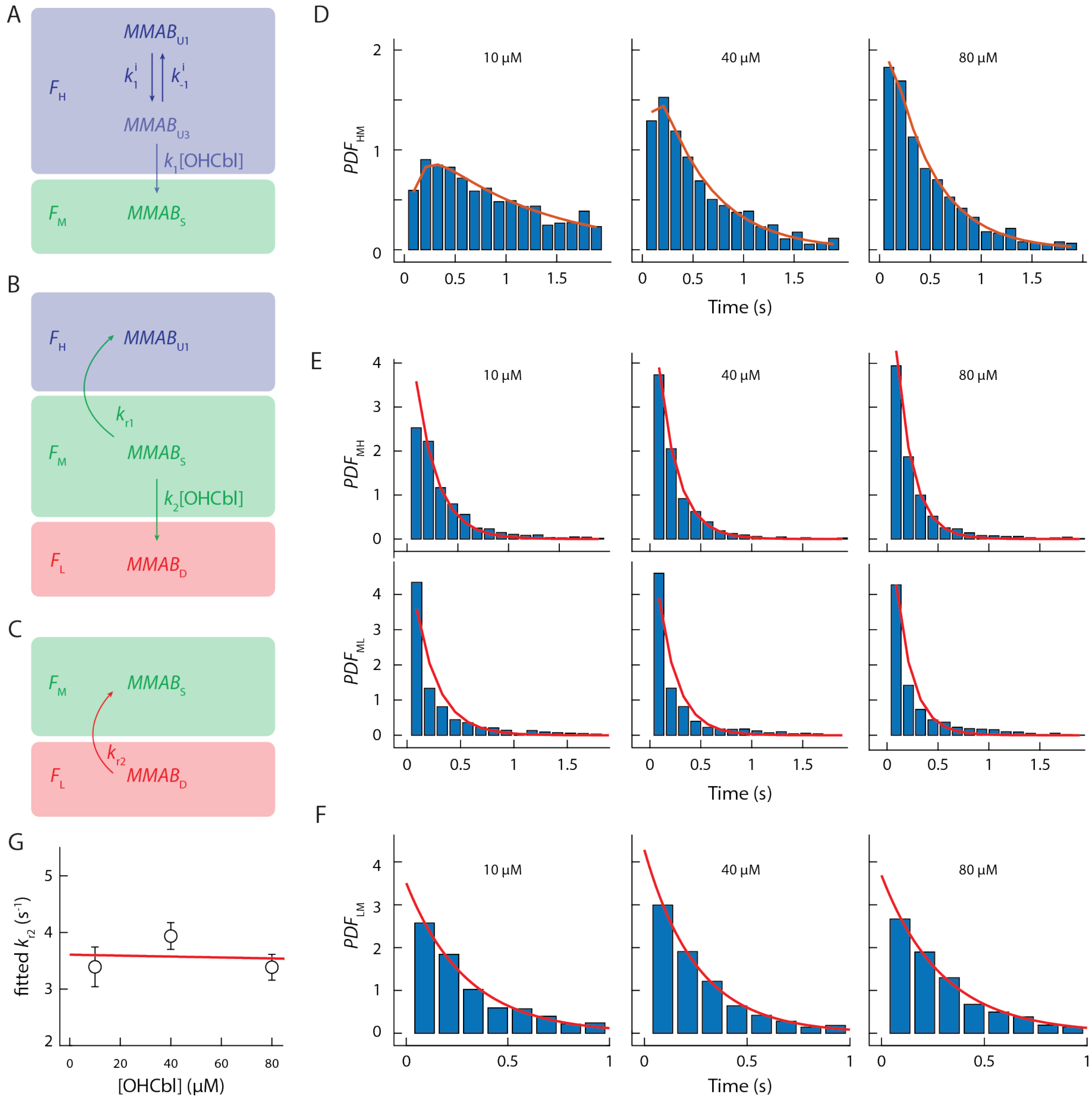


**Supplementary Fig. S11. Extraction of kinetic rate constants for MMAB-OHCbl interactions.** (*A*) Detailed kinetic model for deriving analytical solution of $f_{HM}$. (*B*) Detailed kinetic model for deriving analytical solution of $f_{MH}$ and $f_{ML}$. (*C*) Detailed kinetic model for deriving analytical solution of $f_{LM}$. (*D*) By fitting the experimental dwell time distribution (blue bar) with the analytical solutions (red curve), we extracted rate constants $k_1^i$, $k_{-1}^i$, $k_1$ from $PDF_{HM}$; (*E*) $k_{r1}$ and $k_2$ were extracted from $PDF_{MH}$ and $PDF_{ML}$, (*F*) $k_{r2}$ was extracted from $PDF_{LM}$ (blue bar). (*G*) The $k_{r2}$ is independent of [OHCbl], indicating a single-step dissociation mechanism.

The length of $\tau_{HM}$ (duration time in the $F_H$ state and entering the $F_M$ state) is determined by all the kinetic processes that depart from the $F_H$ state. Based on the submodel of HM (Fig. S11*A*), we derived the analytical expression of the probability density function of dwell time, $f_{HM}(t)$. To solve $f_{HM}(t)$, we started with the analytical expression of the probabilities of finding the species $MMAB_S$, $P_S(t)$. We first focused on the steps that involve the transition $P_{U1}(t)$, $P_{U3}$(t), and $P_S(t)$. The single-molecule rate equations for these steps are:

$$\frac{dP_{U1}(t)}{dt} = k^{i}_{-1}P_{U3}(t) - k^{i}_{1}P_{U1}(t) \qquad S[45]$$

$$\frac{dP_{U3}(t)}{dt} = k^{i}_{1}P_{U1}(t) - (k^{i}_{-1} + k^{o}_{1})P_{U3}(t) \qquad S[46]$$

$$\frac{dP_{S}(t)}{dt} = k^{o}_{1}P_{U3}(t) \qquad S[47]$$

, with the initial condition $P_{U3}(0) = 0$, $P_S(0) = 0$, $P_{U1}(0) = 1$, we obtained the analytical solution of $f_{HM}(t)$:

$$f_{HM}(t) = \frac{dP_{S}(t)}{dt} = k^{o}_{1}P_{U3}(t) = \frac{k^{i}_{1}k^{o}_{1}}{2a}\left[e^{(b+a)t} - e^{(b-a)t}\right] \qquad S[48]$$

, where $a = \sqrt{\frac{1}{4}(k^{i}_{1} + k^{i}_{-1} + k^{o}_{1})^{2} - k^{i}_{1}k^{o}_{1}}$ and $b = -\frac{1}{2}(k^{i}_{1} + k^{i}_{-1} + k^{o}_{1})$.

Using the same principle, we derived the analytical expression of the probability density function of dwell time, $f_{MH}(t)$ and $f_{ML}(t)$ based on submodel Fig. S11*B* with the single-molecule rate equations below:

$$\frac{dP_{U1}(t)}{dt} = k_{r1}P_{S}(t) \qquad S[49]$$

$$\frac{dP_{S}(t)}{dt} = -(k^{o}_{2} + k_{r1})P_{S}(t) \qquad S[50]$$

$$\frac{dP_{D}(t)}{dt} = k^{o}_{2}P_{S}(t) \qquad S[51]$$

, with the initial condition $P_D(0) = 0$, $P_U(0) = 0$, and $P_S(0) = 1$, we get

$$f_{MH}(t) = \frac{dP_{U1}(t)}{dt} = k_{r1}e^{-(k^{o}_{2} + k_{r1})t} \qquad S[52]$$

$$f_{ML}(t) = \frac{dP_{D}(t)}{dt} = k^{o}_{2}e^{-(k^{o}_{2} + k_{r1})t} \qquad S[53]$$

Lastly, we also derived the analytical expression of the probability density function of $f_{LM}(t)$ based on submodel Fig. S11*C* with the single-molecule rate equations and boundary condition below:

$$\frac{dP_{D}(t)}{dt} = -k_{r2}P_{D}(t) \qquad S[54]$$

$$f_{LM}(t) = \frac{dP_{D}(t)}{dt} = k_{r2}e^{-k_{r2}t} \qquad S[55]$$

By globally fitting the $PDF_{HM}$ data of 10, 40, and 80 $\mu$M OHCbl (Fig. S11*D*) with Eq. S48, we got $k^{i}_{1}$ = 2.8 ± 0.3 $s^{-1}$, $k^{i}_{-1}$= 4.0 ± 0.5 $s^{-1}$, $k_1$= 0.28 ± 0.05 $\mu M^{-1}$ $s^{-1}$. Globally fitting $PDF_{MH}$ and $PDF_{ML}$ (Fig. S11*E*) with Eq. S52 and Eq. S53 reports $k_{r1}$(OHCbl) = 4.5 ± 0.6 $s^{-1}$, $k_2$ (OHCbl) = 0.019 ± 0.011 $\mu M^{-1}$ $s^{-1}$. Globally fitting $PDF_{LM}$ (Fig. S11*F*) with Eq. S55 gave the values of $k_{r2}$(OHCbl) = 3.6 ± 0.3 $s^{-1}$.

### 15. Simulation of stopped-flow differential absorption

To compare the extracted rate constants to ensemble results, we simulated the stopped-flow differential absorption by the finite-difference time-domain method. We simulated the time-dependent concentrations of AdoCbl, $MMAB_{U1}$, $MMAB_{U2}$, $MMAB_{S}$ and $MMAB_{D}$ with time step $\Delta t$ = 0.001 s, total concentration of MMAB ($[MMAB]_0$) = 5 µM, and concentrations of AdoCbl ($[\text{AdoCbl}]_0$) varies from 10 to 30 µM. Note that all concentrations mentioned in this section are final concentrations. From the proposed kinetic model, $MMAB_{U1}$ is equilibrated with $MMAB_{U2}$ at $t_0$. This means $k_1^{i}[MMAB_{U1}]_{t_0} = k_{-1}^{i}[MMAB_{U2}]_{t_0}$ and $[MMAB_{U1}]_{t_0} + [MMAB_{U2}]_{t_0} = [MMAB]_0$. With the initial concentrations of $MMAB_{S}$ and $MMAB_{D}$ equal to 0, we derived the changes of each species at any given time, $t_j$, as follows:

$$[MMAB_i]_{t_j} = [MMAB_i]_{t_{j-1}} + \Delta[MMAB_i]_{t_j} \text{ (i = U1, U2, S or D)} \qquad \text{S[56]}$$

$$[\text{AdoCbl}]_{t_j} = [\text{AdoCbl}]_{t_{j-1}} + \Delta[\text{AdoCbl}]_{t_{j-1}} \qquad \text{S[57]}$$

Note that $[\text{AdoCbl}]_{t_j}$ represents the concentration of free AdoCbl at time $t_j$. From the proposed kinetic model in Fig. 3G, we expressed the finite-difference of $[\text{MMAB}_i]$ in $\Delta t$ as shown in Eq S[58] to S[62]:

$$\Delta[MMAB_{U1}]_{t_j} = \Delta t \times (k_{-1}^{i}[MMAB_{U2}]_{t_{j-1}} + k_{r1}[MMAB_S]_{t_{j-1}} - k_1^{i}[MMAB_{U1}]_{t_{j-1}}) \qquad \text{S[58]}$$

$$\Delta[MMAB_{U2}]_{t_j} = \Delta t \times (k_1^{i}[MMAB_{U1}]_{t_{j-1}} - (k_{-1}^{i} + k_1[\text{AdoCbl}]_{t_{j-1}})[MMAB_{U2}]_{t_{j-1}}) \qquad \text{S[59]}$$

$$\Delta[MMAB_S]_{t_j} = \Delta t \times (k_1[\text{AdoCbl}]_{t_{j-1}}[MMAB_{U2}]_{t_{j-1}} + k_{r2}[MMAB_D]_{t_{j-1}} - (k_2[\text{AdoCbl}]_{t_{j-1}} + k_{r1})[MMAB_S]_{t_{j-1}}) \qquad \text{S[60]}$$

$$\Delta[MMAB_D]_{t_j} = \Delta t \times (k_2[\text{AdoCbl}]_{t_{j-1}}[MMAB_S]_{t_{j-1}} - k_{r2}[MMAB_D]_{t_{j-1}}) \qquad \text{S[61]}$$

$$\Delta[\text{AdoCbl}]_{t_j} = \Delta t \times (k_{r1}[MMAB_S]_{t_{j-1}} + k_{r2}[MMAB_D]_{t_{j-1}} - (k_1[MMAB_{U2}]_{t_{j-1}} + k_2[MMAB_S]_{t_{j-1}})[\text{AdoCbl}]_{t_{j-1}}) \qquad \text{S[62]}$$

The absorbance of the solution at 525 nm ($A_{525}$) was estimated from the sum of both bound and free AdoCbl:

$$A_{525} = A_{\text{bound}} + A_{\text{free}} = \varepsilon_{\text{bound}} l[\text{AdoCbl}]_{\text{bound}} + \varepsilon_{\text{free}} l[\text{AdoCbl}]_{\text{free}} \qquad \text{S[63]}$$

, where $\varepsilon$ is the molar absorption coefficient at 525 nm, $l$ is the length of light passing through the sample. Since $[\text{AdoCbl}]_{\text{free}} + [\text{AdoCbl}]_{\text{bound}} = [\text{AdoCbl}]_0$, we substituted $[\text{AdoCbl}]_{\text{bound}}$ with $[\text{AdoCbl}]_0 - [\text{AdoCbl}]_{\text{free}}$ in Eq. S63:

$$A_{525} = l[\varepsilon_{\text{bound}}[\text{AdoCbl}]_0 + (\varepsilon_{\text{free}} - \varepsilon_{\text{bound}})[\text{AdoCbl}]_{\text{free}}] \qquad \text{S[64]}$$

Using $\varepsilon_{\text{bound, 525m}}$ = 0.0081 $\mu M^{-1}cm^{-1}$ and $(\varepsilon_{\text{free}} - \varepsilon_{\text{bound}})$ = $6.69\times10^{-3} \mu M^{-1}cm^{-1}$ , we simulated the time-dependent absorption trajectories with time step $\Delta t$ = 0.001 s, total MMAB concentration ($[\text{MMAB}]_0$) = 5 µM and $[\text{AdoCbl}]_0$ varies from 10 to 30 µM. The simulation results show the same trend as Banerjee's stopped-flow absorption data (7) as shown in Fig. S12.

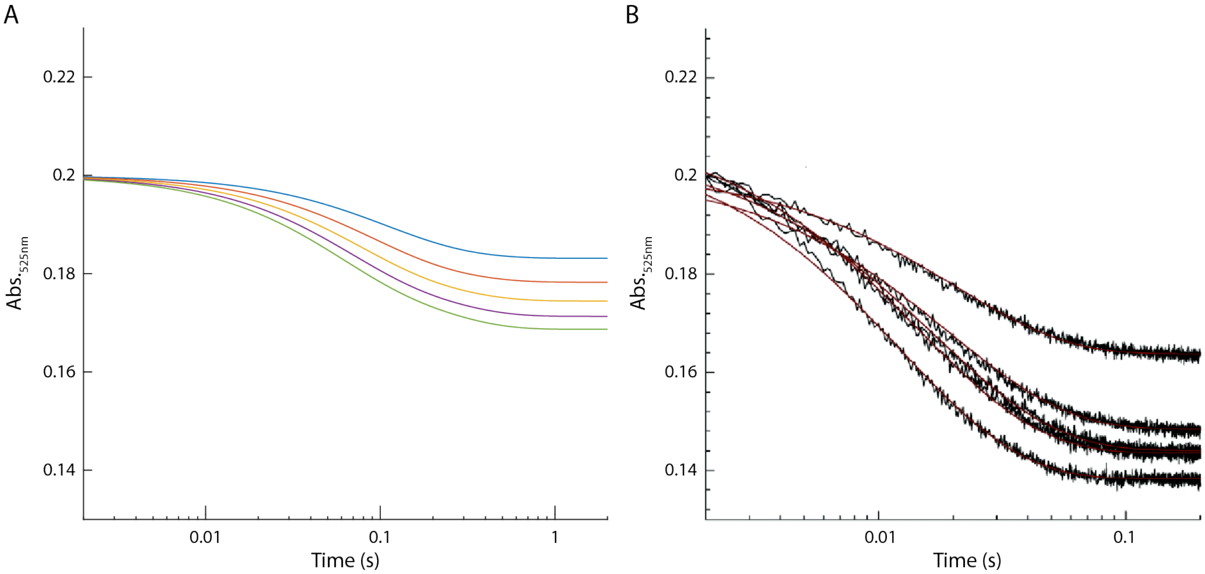


**Supplementary Fig. S12. Comparison of stopped-flow absorption results of MMAB.** (*A*) Stopped-flow absorption simulation using kinetic model and rate constants determined in this work. (*B*) ensemble experimental data adapted from ref. S7.
